# Simvastatin attenuates disease phenotypes in human induced pluripotent stem cell models of familial Parkinson’s disease through RhoA inhibition

**DOI:** 10.64898/2026.08.26.747232

**Authors:** Sissel Ida Schmidt, Justyna Okarmus, Matias Ryding, Ida Keller Skousen, Nicolaj Frederik Broner Jensen, Emil Birch Christensen, Lucas S. Winkelmann, Alice Dupont Juhl, Miamaja Klæbel, Morten Blaabjerg, Kristine Freude, Daniel Wüstner, Richard Wade-Martins, Brent Ryan, Morten Meyer

## Abstract

**Background:** Statins have gained increasing interest for their potential therapeutic effect in Parkinson’s disease (PD). Beyond their cholesterol-lowering effect, statins decrease synthesis of isoprenoids, which is believed to account for their pleiotropic effects. Isoprenylation is important for proper membrane localization and function of the Rho GTPases, including RhoA. RhoA signalling has emerged as a possible underlying signalling pathway involved in the pathogenesis of PD and other neurodegenerative diseases.

**Methods:** In the present study, we investigated the effects of simvastatin on neurodegeneration-associated phenotypes using human induced pluripotent stem cell-derived dopaminergic (DA) neurons from both PD patients and isogenic *PARK2^-/-^* cell lines. The dependence on RhoA was confirmed using direct RhoA inhibition using rhosin. Assessed phenotypes included structural integrity, mitochondrial and lysosomal characteristics, cytokine secretion, and cell viability. To understand the relevance of RhoA in PD, RhoA activity was measured in 32 PD patient iPSC-derived lines with different familial PD-related mutations and in healthy controls.

**Results:** Simvastatin rescued multiple PD-associated phenotypes, including impaired DA neurite outgrowth, mitochondrial and lysosomal alterations, cytokine release, and cell death. RhoA inhibition was associated with changes in mitophagy- and autophagy-related markers, suggesting improved autophagic and mitophagic turnover. Furthermore, we performed the first systematic screen of RhoA activity across 32 iPSC-derived DA neuron lines representing multiple genetic forms of PD (PINK1 loss of function, parkin loss of function, *LRRK2* (G2019S), *LRRK2* (R1441C), *GBA* (L44P), *GBA* (N370S), *A53T,* and *SNCA* triplication) and healthy controls. RhoA activity was perturbated across several genetic forms of PD subtypes and was significantly increased in many, although not all, patient lines compared with healthy controls, highlighting disease heterogeneity and supporting RhoA dysregulation as a shared pathogenic mechanism in a subset of PD.

**Conclusions:** Our findings identify aberrant RhoA signalling as a convergent pathogenic mechanism across multiple forms of genetic PD and demonstrate that simvastatin ameliorates PD-associated phenotypes through RhoA inhibition. These results support RhoA as a promising therapeutic target while emphasizing the importance of patient stratification based on RhoA activity.

## BACKGROUND

Parkinson’s disease (PD) is the second most common neurodegenerative disease and is caused by the progressive death of dopaminergic (DA) neurons in the substantia nigra pars compacta (SNpc) of the midbrain^1, 2^. Although the etiology of PD remains inconclusive, studies of gene mutations linked to PD, including *SNCA* (α-synuclein), *PARK2* (parkin), *PINK1* (PTEN-induced putative kinase 1), *LRRK2* (Leucine-rich repeat serine/threonine-protein kinase 2), and *GBA* (glucocerebrosidase), have identified several cellular mechanisms involved in the DA neuronal degeneration. These include dysfunctional mitochondrial homeostasis, increased oxidative stress, protein misfolding and impairments in the ubiquitin-proteasome and autophagy-lysosomal systems, and neuroinflammation^3–5^.

Ras homolog gene family member A (RhoA) signalling has recently emerged as a possible underlying signalling pathway involved in the pathogenesis of PD and other neurodegenerative diseases^6, 7^. RhoA is a small GTPase of the Rho family that regulates several cellular processes such as cytoskeletal rearrangement, cell migration, and cell death^6, 8, 9^. In mice treated with the DA neurotoxin 1-methyl-4-phenyl-1,2,3,6-tetrahydropyridine (MPTP), RhoA and its downstream effector protein ROCK were found upregulated in the SNpc, and ROCK inhibition was reported to protect against the MPTP-induced DA cell death^10–12^. Similarly, MPTP and the neurotoxic pesticide rotenone are reported to increase RhoA signalling in cell models of PD, and inhibition of RhoA signalling ameliorated the neurotoxin-induced DA axonal degeneration^13–15^. While the role of RhoA signalling is well established in neurite retraction, spine and synapse loss, and neuronal apoptosis, much less is known about the role of RhoA signalling in mitochondrial homeostasis and autophagy, two important players in PD.

Loss-of-function mutations in the *PARK2* gene lead to impaired mitochondrial quality control and cause autosomal recessive PD with early disease onset^16^. *PARK2* encodes the ubiquitin E3 ligase parkin, which is recruited to depolarized mitochondria by the mitochondrial serine/threonine kinase PINK1 and ubiquitinates proteins on the outer mitochondrial membrane (OMM), thus targeting the damaged mitochondria for degradation by mitophagy^17^.

We have previously compared *PARK2*^-/-^ and isogenic control induced pluripotent stem cell (iPSC)-derived neurons using a novel, large-scale mass spectrometry-based proteomics and post-translational modification (PTM)-omics approach, in order to map changes in protein profiles and PTMs caused by parkin deficiency in neurons (dataset available through the ProteomeXchange Consortium, identifier: PXD008894)^18^. Pathway analysis and *in vitro* assays revealed altered migration and impaired neurite outgrowth in *PARK2^-/-^* neurons. RhoA signalling was identified as a key upstream regulator, and RhoA activity was significantly increased in *PARK2^-/-^* neurons^18^.

Interestingly, the lipophilic statin *simvastatin* has been found to inhibit RhoA in the brain^19, 20^. Statins inhibit 3-hydroxy-3-methylglutaryl-coenzyme A (HMG-CoA) reductase, the rate-limiting enzyme in cholesterol biosynthesis, and are widely used in the clinic for treating hypercholesterolemia^21^. Besides their cholesterol-lowering effect, statins decrease synthesis of isoprenoids, such as farnesyl pyrophosphate (FPP) and geranylgeranyl pyrophosphate (GGPP), which might explain some of the pleiotropic effects of statins^19^. Isoprenylation is important for proper membrane localization and function of the Rho GTPases, including RhoA, Rac, and Cdc42, and depletion of isoprenoids by statins results in accumulation of non-functional Rho GTPases in the cytoplasm^22, 23^. Hence, simvastatin has been suggested to exert neuroprotective effects.

In the present study, we investigated whether a pharmacologically relevant concentration of simvastatin (300 nM)^24^ exerts neuroprotective effects in human iPSC-derived DA neurons modelling PD. Using both isogenic *PARK2^-/-^*and patient-derived PD models, we examined the effects of simvastatin on key PD-associated cellular phenotypes, including neuronal migration, neurite outgrowth, mitochondrial and lysosomal homeostasis, inflammatory responses, and cell survival. To determine whether these effects are mediated through RhoA signalling, we compared simvastatin treatment with direct pharmacological RhoA inhibition. Finally, to investigate the relevance of RhoA signalling across genetic forms of PD, we profiled RhoA activity in a panel of patient-derived iPSC DA neurons carrying mutations in *PINK1*, *PARK2*, *LRRK2*, *GBA*, and *SNCA*.

## METHODS

### Neural stem cell (NSC) propagation and differentiation

*PARK2*^-/-^ and isogenic control iPSC-derived NSCs were obtained from XCell Science Inc. (Novato, CA, USA, cell line info in Table S1). Cell line characterization at both iPSC and NSC stage are previously described^18, 25^.

NSCs were propagated on Geltrex (Gibco) coated plates in Neurobasal medium (Gibco) supplemented with 1% Non-Essential Amino Acids (NEAA, Gibco), 1% GlutaMAX-1 (Gibco), 1% B27 (Gibco), 1% penicillin-streptomycin (Sigma), and 10 ng/ml basic fibroblast growth factor (bFGF, Sigma). The medium was changed every second day, and NSCs were passaged 1:3 when 80-90% confluent, by dissociating the cells for 5 min with accutase (Gibco).

NSCs were differentiated using a commercially available dopaminergic differentiation kit (XCell Science) for at least 25 days (Fig. 1A). In short, NSCs were seeded onto 20 μg/ml poly-L-ornithine (Sigma) and 10 μg/ml laminin (Thermo Fisher) coated plates at a density of 50,000 cells/cm^2^. The first 9 days of differentiation were carried out in DOPA Induction Medium (XCell Science) supplemented with DOPA Induction Supplement A, B and C (XCell Science) and 200 ng/ml rh-sonic hedgehog (SHH, Peprotech) with medium change every second day. At day 5 and 10, cells were passaged using accutase and replated at a desired cell density depending on the final experiment. At day 10, medium was switched to DOPA Maturation Medium (XCell Science) supplemented with DOPA Maturation Supplement A (XCell Science) until day 16, when the maturation medium was supplemented with DOPA Maturation Supplement B (XCell Science) until at least day 25 with 50% medium change every second day.

**Figure 1.**
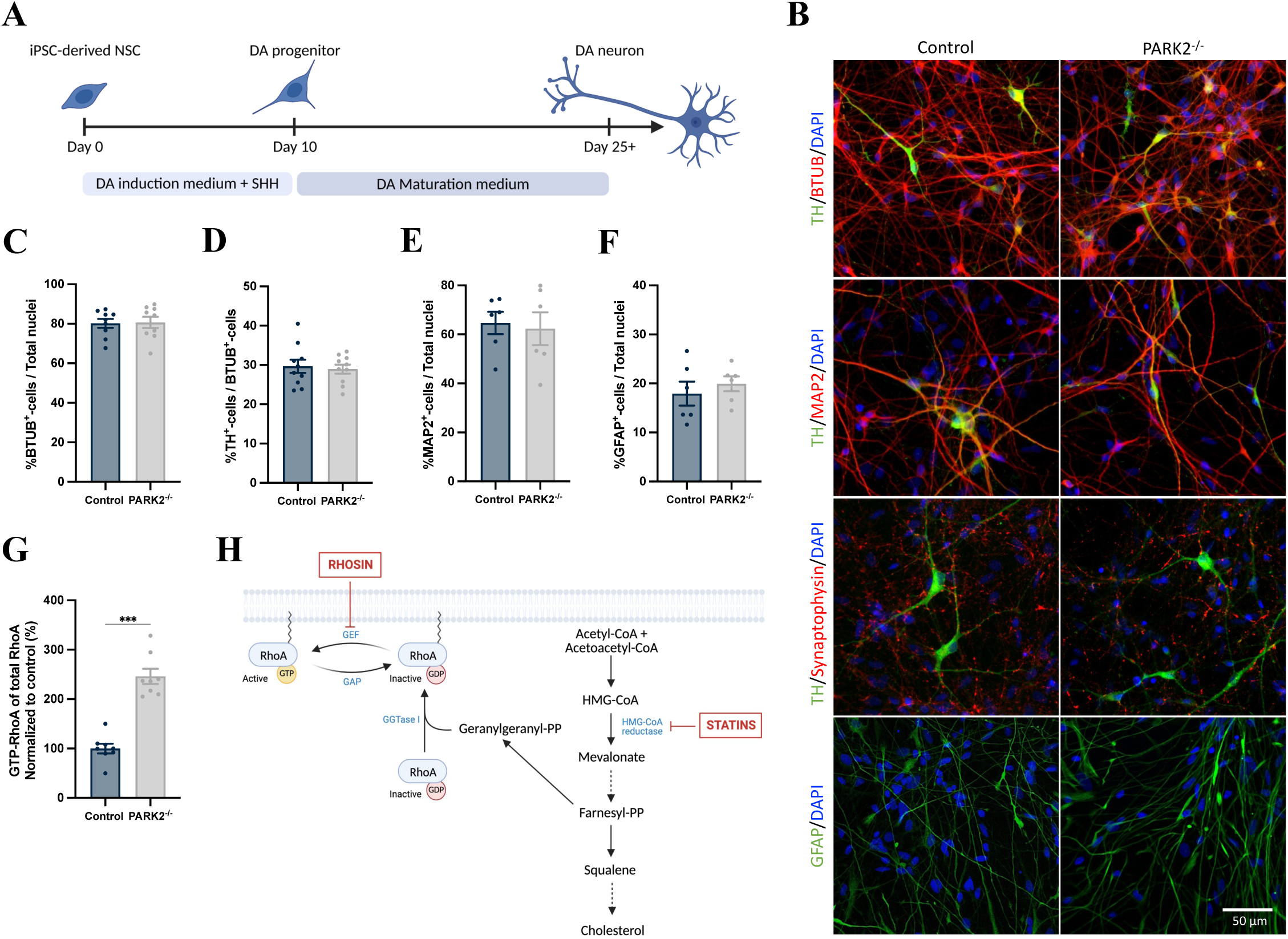
Increased RhoA activity in *PARK2*^-/-^ neurons. A) Graphical overview of the applied commercially available dopaminergic (DA) differentiation kit for differentiating iPSC-derived *PARK2*^-/-^ NSCs and isogenic control NSCs. B-F) Immunofluorescence staining and quantification of β-tubulin-III^+^ (BTUB, red), tyrosine hydroxylase (TH, green), microtubule-associated protein 2 (MAP2, red), synaptophysin (red), and glial-acidic fibrillary protein (GFAP, green) of differentiated cultures showed no difference in the differentiation capacity of *PARK2*^-/-^ and isogenic control cells or maturity of the neuronal cultures. Scale bar = 50 µm. Mean ± SEM, n = 6-9 coverslips per cell line from 2 independent differentiations. Statistical analysis: Student’s t-test. G) RhoA activity was significantly increased in *PARK2*^-/-^ neurons, measured from the ratio of GTP-bound RhoA to total RhoA levels. Mean ± SEM, n = 8 samples per cell line from 4 independent differentiations. Statistical analysis: Student’s t-test, ***p < 0.001. H) Graphical overview of the RhoA-inhibiting strategies applied in this study. RhoA activity was either inhibited by blocking GTP binding using rhosin (120 µM) or by preventing isoprenylation of RhoA by simvastatin (300 nM).

Cells were treated with 120 µM rhosin (Tocris), 300 nM simvastatin (Sigma) or 50 µM gemfibrozil (Sigma) from day 18 until day 25, if not otherwise specified, and replenished every second day with medium change. Inhibition verification and dose-response and toxicity data are found in the Supplementary Information (Fig. S2-S4).

### PD patient iPSC generation, propagation, and differentiation

iPSC lines were generated from skin biopsies from PD patients and controls recruited to the Discovery clinical cohort established by the Oxford Parkinson’s Disease Center. The PD patients fulfilled the UK Brain Bank diagnostic criteria for clinically probable PD at presentation. Participants gave signed informed consent to mutation screening and derivation of iPSC lines (Ethics committee: National Health Service, Health Research Authority, NRES Committee South Central, Berkshire, UK, REC 10/H0505/71). Fibroblasts from skin biopsies were reprogrammed to iPSC lines as previously described^26^. This study included iPSC lines from three PD patients with *PARK2*^27, 28^ mutations, five with *PINK1*^27,28^ mutations, four with *LRRK2*-*G2019S*^29^ mutation, three with *LRRK2-R1441C* ^30^ mutation, four with *GBA-L444P* mutation, two with *GBA-N370S*^31^, two with *SNCA-A53T*^32^, three with *SNCA-triplication* ^32^ and five healthy controls (cell line info in Table S1).

iPSCs were propagated as a monolayer on geltrex-coated plates in complete mTeSR1 medium (Stem Cell Tech.) with 1% penicillin-streptomycin and passaged 1:2-3 when 80% confluent using TrypLE (Gibco). Medium change was performed every day. ROCK inhibitor (Y27632, Bio-Techne) was added to the medium to promote survival when thawing or passaging the iPSCs.

iPSCs were differentiated using a modified dual-SMAD inhibition protocol as previously described^29^ (Fig. 6A). In brief, iPSCs were plated at a density of 150,000 cells/cm^2^ on Geltrex and patterned for 11 days to become ventral midbrain precursors in knockout serum replacement medium (KSR) containing KO DMEM (Life Technologies), 15% knockout serum replacement (Life Technologies), 2 mM L-glutamine (Life Technologies), 10 mM β-mercaptoethanol (Sigma), and 1% NEAA supplemented with 118 nM LDN193189 (Sigma), 10 µM SB431542 (Tocris, day 0-4), 100 ng/ml SHH-C24II (Bio-Techne, day 1-6), 2 µM Purmorphamine (Millipore, day 1-6), 100 ng/ml FGF8a (Bio-Techne, day 1-6), and 3 µM CHIR99021 (Tocris, from day 3) with medium change every day. From day 5 of differentiation, the KSR medium was gradually shifted to NNB medium containing Neurobasal medium, 0.5X N2 (Gibco), 0.5X B27, and 2 mM L-glutamine. Cells were expanded at the day 11 stage for 3 weeks in KSR medium mixed with NNB medium 1:4 and supplemented with 118 nM LDN193189 and 3 µM CHIR99021. Following expansion, ventral midbrain precursor cells were matured to dopaminergic neurons in NB medium containing Neurobasal medium, 1% B27, and 2 mM L-glutamine supplemented with 3 µM CHIR99021 (until day 13), 20 ng/ml brain-derived neurotrophic factor (BDNF, Peprotech), 20 ng/ml glial cell line-derived neurotrophic factor (GDNF, Peprotech), 1 ng/ml transforming growth factor type β3 (TGFβ3, Peprotech), 10 µM DAPT (Abcam), 0.2 mM ascorbic acid (Sigma), and 0.5 mM dibutyryl cAMP (Sigma). At day 20, cells were passaged using accutase and replated on 0.1 mg/ml PLO (Sigma) and 10 µg/ml Biolaminin (BioLamina) coated plates at a desired cell density depending on the final experiment. 1X Antibiotic-Antimycotic (Sigma) was added to the medium from day 20. On day 22, neuronal cultures were treated with 1 µg/ml Mitomycin C (Life Technologies) in NB medium for 1 hour to reduce the number of undifferentiated proliferative cells in the cultures and matured for an additional two weeks with medium change every second day until day 35 before any given experiment.

Treatment with 300 nM simvastatin was initiated at day 30 and replenished every second day with medium change until day 35. Dose-response and toxicity data are found in the Supplementary Information (Fig. S6).

### Immunofluorescence staining

Neuronal cultures were fixed in 4% paraformaldehyde (PFA, Sigma) in 0.1 M phosphate-buffered saline (PBS, pH 7.4) for 15 min at room temperature (RT) and washed twice in PBS for 15 min before permeabilization and blocking of unspecific antibody binding in either 0.05 M TBS/0.1% Triton X-100 (Sigma)/5% goat or donkey serum (Millipore) or 0.05 M TBS/0.1% saponin (Sigma)/5% goat or donkey serum blocking buffer, depending on the staining, for 1 hour at RT. Primary antibodies were diluted in blocking buffer and incubated overnight (ON) at 4°C. The following primary antibodies and dilutions were applied: rabbit anti-tyrosine hydroxylase (TH, Millipore) 1:600, sheep anti-TH (Millipore), goat anti-FOXA2 (Bio-Techne) 1:250, rabbit anti-LMX1A (Abcam) 1:250, rabbit anti-β-tubulin III (Sigma) 1:2000, mouse anti-MAP2a+b (Sigma) 1:2000, chicken anti-MAP2a+b+c (Millipore) 1:2000, mouse anti-synaptophysin (Sigma) 1: 200, guinea pig anti-synapsin (Synaptic systems) 1:250, Rabbit anti-homer 1 (Synaptic systems) 1:250, rabbit anti-glial fibrillary acidic protein (GFAP, DAKO) 1:4000, rabbit anti-TOM20 (Santa-Cruz) 1:1000, mouse anti-TOM20 (Santa-Cruz) 1:500, rabbit anti-LAMP1 (Abcam) 1:1000, mouse anti-LAMP1 (Santa-Cruz) 1:250, rabbit anti-cleaved caspase 3 (Cell Signalling) 1: 400, rabbit anti-NIX (Abcam) 1:1000.

Cultures were washed three times in either 0.05 M TBS/0.1% Triton X or 0.05 M TBS/0.1% saponin wash solution for 15 min at RT and incubated with fluorophore-conjugated secondary antibodies, diluted in blocking buffer, for 2 hours at RT. The following secondary antibodies were applied in dilution 1:500: goat anti-rabbit AF488 (Thermo Fisher), goat anti-mouse AF488 (Thermo Fisher), goat anti-guinea pig AF488 (Abcam), donkey anti-rabbit AF488 (Thermo Fisher), donkey anti-mouse AF488 (Thermo Fisher), donkey anti-goat AF488 (Thermo Fisher), donkey anti-sheep AF488 (Thermo Fisher), goat anti-rabbit AF555 (Thermo Fisher), goat anti-mouse AF555 (Thermo Fisher), donkey anti-rabbit AF555 (Thermo Fisher), donkey anti-mouse AF555 (Thermo Fisher), goat anti-chicken AF647 (Thermo Fisher), donkey anti-rabbit AF647 (Thermo Fisher), donkey anti-sheep AF647 (Thermo Fisher).

Cultures were washed twice in 0.05 M TBS for 15 min at RT and cell nuclei were counterstained with 10 µM 4",6-diamidino-2-phenylindole dihydrochloride (DAPI, Sigma). Cultures grown on coverslips were afterwards mounted on glass slides using ProLong® Diamond mounting medium (Molecular Probes). For cultures imaged directly in 96- or 384-well plates, DAPI solution was aspirated and washed once before left in PBS for imaging. Fluorescence images were acquired on either the FluoView FV1000MPE multiphoton laser confocal microscope (Olympus), BX53 fluorescence microscope (Olympus), the ImageXpress Automated Imaging System (Molecular Device), or the Opera Phenix High-Content Screening confocal microscope (Perkin-Elmer) at 20X or 63X magnification in a blinded manner on at least five randomly chosen fields per coverslip/well from independent differentiations.

Cell counts for general characterization of *PARK2*^-/-^ and isogenic control cultures were quantified manually in ImageJ using the Cell Counter Plugin. Only cells displaying an extensive immunostaining with distinct cellular morphology were counted. Mitochondrial and lysosomal area were analysed automated in ImageJ by converting images to binary format and using the Analyse Particles function. All staining quantifications were normalized to total cell numbers as evaluated by automated quantification of DAPI^+^ nuclei in ImageJ.

All image analysis of patient-derived cultures was quantified automated using the Harmony imaging software (Perkin-Elmer) and normalized to total cell counts.

### Western blotting

Cell pellets were lysed in RIPA buffer (Thermo Fisher) with protease (Complete tablets, Roche) and phosphatase inhibitor (PhosSTOP tablets, Roche) and sonicated for 2 x 10 sec at amplitude 2 microns on ice. Protein concentrations were determined using the bicinchoninic acid assay (BCA, Pierce), and equal amounts of protein were denatured for 10 min at 70°C before being loaded on 4-12% Bis-Tris gels (Thermo Fisher) together with SeeBlue™ Plus2 Pre-stained Protein Standard (Thermo Fisher) for molecular weight estimations. Proteins were separated at 200 V for 50 min in MOPS running buffer (Thermo Fisher) with 0.25% antioxidant (Thermo Fisher) and transferred to polyvinylidene difluoride (PVDF) membranes (Invitrogen) at 20V for 8 min using the iBlot transfer system (Invitrogen). Membranes were blocked for 60 min at RT in 0.05M TBS/0.05% Tween-20 (Sigma)/5% skim milk (Natur Drogeriet)/5% BSA (Sigma) (blocking buffer) before ON incubation at 4°C with primary antibodies diluted in blocking buffer. The following primary antibodies and dilutions were applied: mouse anti-Parkin (Cell Signalling) 1:1000, rabbit anti-NIX (Cell Signalling) 1:1000, mouse anti-P62/SQSTM1 (Cell Signalling) 1:500, rabbit anti-LC3B (Cell Signalling) 1:1000, and mouse anti-actin (Millipore) 1:6000. Membranes were washed in 0.05 M TBS/0.05% Tween-20 and incubated for 1 hour at RT with horseradish peroxidase (HRP)-conjugated anti-mouse or -rabbit IgG (DAKO) diluted 1:2000 in blocking buffer. Membranes were then repeatedly washed in 0.05 M TBS/0.05% Tween-20, and protein expression was detected using ECL (Thermo Fisher) with the ChemiDoc MP imaging system (BioRad) and quantified by densitometry using Image Lab software (BioRad).

### Membrane fractionation and Western blotting in NSCs

To assess RhoA distribution in total and membrane fractions, PARK2^-/-^ and isogenic control NSCs were cultured to confluency and treated for 24 hours with vehicle (DMSO), 300 nM simvastatin, 5 µM geranylgeranyl transferase inhibitor (GGTI), or 5 µM farnesyl transferase inhibitor (FTI). Control cultures received DMSO only. Following treatment, cells were harvested and fractionated using the Mem-PER™ Plus Membrane Protein Extraction Kit (Thermo Fisher Scientific) according to the manufacturer’s instructions to obtain membrane protein fractions. Total protein lysates were prepared in parallel. Due to low protein yield in membrane fractions, samples were concentrated using Vivaspin 500 centrifugal concentrators (Sartorius) prior to protein quantification. Protein concentrations were determined using the bicinchoninic acid (BCA) assay as described above. Equal amounts of protein (15 µg) were subjected to SDS-PAGE and Western blotting according to the protocol described above. RhoA protein levels were detected using mouse anti-RhoA (Santa Cruz) 1:400. Membrane fractions were normalized to rabbit anti-ATPase^a2^ (Millipore) 1:5000, and total lysates were normalized to mouse anti-actin (Millipore) 1:6000. Band intensities were quantified by densitometry using Image Lab software (BioRad).

### RhoA activity assay

The ratio of active GTP-bound RhoA to total RhoA was measured using the G-LISA RhoA Activation Assay Biochem Kit (Cytoskeleton Inc.) and Total RhoA ELISA Kit (Cytoskeleton Inc.) according to the manufacturer’s protocol.

### Migration assay

A scratch assay was used to evaluate cell migration of *PARK2*^-/-^ and isogenic control neurons. Cells were plated in a 96-well plate at day 10 of differentiation, and at day 19 a scratch was performed in the centre of each well using an Incucyte Wound Maker 96-Tool (Sartorius) followed by a full medium change to remove cell debris and initiate treatment. The scratch area was bright-field imaged at day 18 and again at day 24 using the Eclipse TS100 inverted microscope (Nikon). The distance of migrating cell bodies into the scratch area from day 18 to day 24 was quantified using ImageJ with the investigator blinded.

### Neurite outgrowth assays

Neurite outgrowth was assessed in *PARK2*^-/-^ and isogenic control cells. For baseline neurite measurements, cells were seeded at low cell density (20.000 cells/cm^2^) at day 10 of differentiation. Where indicated, pharmacological treatments were applied at day 10 and maintained for three days. These included vehicle (DMSO), rhosin, 200 nM simvastatin, 5 µM geranylgeranyl transferase inhibitor (GGTI), or 5 µM farnesyl transferase inhibitor (FTI). At day 13, cells were fixated in 4% paraformaldehyde and immunolabelled for TH and DAPI using the standard staining protocol described above. Imaging was performed using a BX53 fluorescence microscope (Olympus), and average neurite length/cell was measured using the NeuronJ plugin for ImageJ. Only cells with intact soma and one or more neurites were included, whereas overlapping neurites were excluded from the analysis.

For parkin PD patient iPSC-derived dopaminergic neurons, the scratch assay was used to assess neurite outgrowth into the scratch area (scratch recovery). Cells were plated in 96-well plates at day 20 of differentiation, and scratch was performed at day 30 in the centre of each well using a 10 µl pipette tip, followed by a full medium change to remove cell debris and initiate treatment. The scratch area was imaged every second day using the Opera Phenix High-Content Screening Confocal microscope (PerkinElmer). The density of the scratch area covered by processes for each day were analysed using ImageJ. Values from day 30 were subtracted as background, and data were adjusted for differences in cell density and scratch area size.

### ATP assay

The level of intracellular ATP was measured in *PARK2*^-/-^ and isogenic control cultures using the ATPlite Assay (Perkin Elmer) according to the manufacturer’s instructions. The luminescence signal was recorded using an Orion L Microplate Luminometer (Titertek Berthold).

### β-galactosidase activity assay

β-galactosidase activity was measured using the Fluorometric β-Galactosidase Activity Assay Kit (BioVision) according to the manufacturer’s protocol. The fluorescence signal was recorded in kinetic mode using a Paradigm multi-mode microplate reader (Beckman Coulter), and β-galactosidase activity was calculated from two timepoints in the linear range. Assay values were normalized to sample protein concentrations measured by the BCA.

### Cell death assays

Necrotic cell death was evaluated by measuring the release of lactate dehydrogenase to conditioned culture medium using the CytoTox96® Non-Radioactive Cytotoxicity Assay (Promega) according to the manufacturer’s instructions. Cell death was evaluated for different timepoints and treatment durations, as shown schematically in Fig. 5A. Data were normalized to total cell counts for the corresponding well that the conditioned culture medium was collected from.

Apoptotic cell death was evaluated by immunostaining for cleaved caspase 3 according to the method described for immunocytochemistry.

### Cytokine and neurofilament light chain release

Release of cytokines and neurofilament light chain into the culture medium was measured using the V-PLEX Proinflammatory Panel 1 Mouse kit (MesoScale Discovery) and the R-PLEX Human Neurofilament L Assay (MesoScale Discovery), respectively. Conditioned culture medium was collected at day 45 of differentiation with treatment initiated at day 18, and assays were performed according to the manufacturer’s instructions. Data were normalized to total cell counts for the corresponding well that the conditioned culture medium was collected from.

### Mitochondrial membrane potential

To measure mitochondrial membrane potential (Δψm), the JC-10 assay was performed (AAT Bioquest, Inc.). JC-10 is a cationic, lipophilic dye that forms reversible red-fluorescent aggregates in polarized mitochondria. Impaired mitochondrial membrane potential causes JC-10 to leak out of the mitochondria and return to its monomeric green-fluorescent form.

Cells were washed with neurobasal medium w/o phenol red before being loaded with 3 µM JC-10 diluted in NNB medium w/o phenol red and incubated at 37°C for 1 hour. Following incubation, cells were washed once in neurobasal medium w/o phenol red and placed in NNB medium w/o phenol red for reading. The fluorescence intensities were measured using the PHERAstar FSX plate reader (BMG Labtech) by fluorescence excitation/emission maxima: 485/520 nm (monomer form) and 540/590 nm (aggregate form). To confirm that the JC-10 signal was indicative of Δψm, experiments were terminated by inducing maximal mitochondrial depolarization by addition of 20 µM carbonyl cyanide 3-chlorophenylhydrazone (CCCP, Sigma). The ratio of red/green florescence (ΔF, 590 nm/520 nm) was calculated before and after addition of CCCP, and the relative fluorescence ratio (ΔF-ΔF_CCCP_/ΔF) used to measure Δψm.

### Statistical analysis

All statistical analyses and graphical representations were performed in Prism version 9 (GraphPad software). Data are presented as mean ± SEM, and the specific n-values and statistical tests used are noted in each figure legend. Datasets were tested for normality using the D’Agostino-Pearson test. Two-tailed unpaired Student’s t test was used to compare the means of two groups. One-way analysis of variance (ANOVA) with Dunnett’s multiple comparison test was used to compare the means of more than two groups, and two-way ANOVA with Turkey multiple comparison test was used to compare the means across grouped data.

In the case of too low a sample size for normality testing, normal distribution of the dataset was assumed. Statistical p-values < 0.05 were considered significant, and specific p-values are denoted by asterisks on the figure graphs as follows: NS = not significant, *p < 0.05, **p < 0.01, ***p < 0.001.

## RESULTS

### Increased RhoA activity in *PARK2*^-/-^ neurons

To study the effect of RhoA inhibition in PD, we utilized *PARK2^-/-^* and isogenic control iPSC-derived neurons with previously established increased RhoA activity^18^. The *PARK2^-/-^* iPSC line was created from a healthy control iPSC line by zinc finger nuclease gene editing as previously described and characterized ^18, 25, 33, 34^. *PARK2^-/-^*and isogenic control iPSC-derived neural stem cells (NSCs) were differentiated for at least 25 days (Fig. 1A) into neuronal cultures expressing a large percentage of β-tubulin-III^+^ neurons (>80%), of which >30% were tyrosine hydroxylase (TH)-expressing DA neurons (Fig. 1B-D). Midbrain identity of the DA neurons has previously been confirmed by qPCR for the midbrain DA neuron-specific markers EN1, LMX1A, NURR1 and GIRK2^33, 34^ and staining for FOXA2^27^. The maturity of the cultures was verified by robust expression of microtubule-associated protein 2 (MAP2) (65%) and the presynaptic marker synaptophysin (Fig. 1B and E). This was further supported by calcium imaging recordings indicating neuronal functionality as the neurons displayed spontaneous activity and could be depolarized by potassium chloride (Fig. S1). The cultures also contain GABAergic neurons^18^ and glial fibrillary acidic protein (GFAP)^+^ astrocytes (Fig. 1B and F). Both cell lines showed comparable expression levels for all markers with no significant difference, demonstrating equal differentiation capacity of the two cell lines. Despite comparable proportions of GFAP^+^ astrocytes in *PARK2^-/-^*and isogenic control cultures, in line with previous observations, qualitative differences in GFAP staining intensity and astrocytic morphology were observed, with *PARK2^-/-^* cultures displaying more pronounced staining and hypertrophic-like features^18, 35^.

Increased RhoA activity was also observed in *PARK2^-/-^*neuronal cultures measured by the ratio of active GTP-bound RhoA to total RhoA (Fig. 1G). RhoA cycles between an inactive guanosine diphosphate (GDP)-bound state in the cytosol and an active guanosine triphosphate (GTP)-bound state associated with the plasma membrane^6^. In the present study, we apply two different strategies for inhibiting RhoA: either by directly blocking the GTP binding to RhoA using the compound rhosin^36^ or by inhibiting the isoprenylation of RhoA, required for its membrane association, using the lipophilic statin simvastatin^19^ (Fig. 1H). Inhibition verification is found in the Supplementary Information (Fig. S2).

### RhoA inhibition rescues perturbations in migration and neurite outgrowth of *PARK2*^-/-^ neurons

RhoA signalling is known to promote neuroblast migration and inhibit neuritogenesis^6, 37^, phenotypes that we have previously established for *PARK2^-/-^* neurons^18^. To test the effect of simvastatin treatment on migration of the *PARK2^-/-^* neurons, we performed a scratch assay on day 18 of differentiation and evaluated the migration of cell bodies into the scratch area after 7 days. In accordance with our previous findings, *PARK2^-/-^* neuroblasts migrate significantly more into the scratch area than healthy isogenic controls (Fig. 2A-B). Both treatment with simvastatin and direct RhoA inhibition using rhosin for the duration of the scratch assay were found to normalize the migration completely to levels equal to untreated control (Fig. 2A-B, for dose response see Fig. S3 and S4). Treatment with the fibrate gemfibrozil, which reduces cholesterol levels without having a RhoA-inhibiting effect^38^, did not rescue the increased migration for any of the concentrations tested (Fig. S5A-B). Moreover, neither gemfibrozil nor simvastatin treatment was found to reduce intracellular cholesterol levels in the *PARK2^-/-^* or control cultures as evaluated by filipin staining (Fig. S5C-D). This is in accordance with previous studies showing that cholesterol-free medium is necessary to decrease cellular levels of cholesterol with statin treatment, as cholesterol can otherwise be replenished from the culture medium^39, 40^.

**Figure 2.**
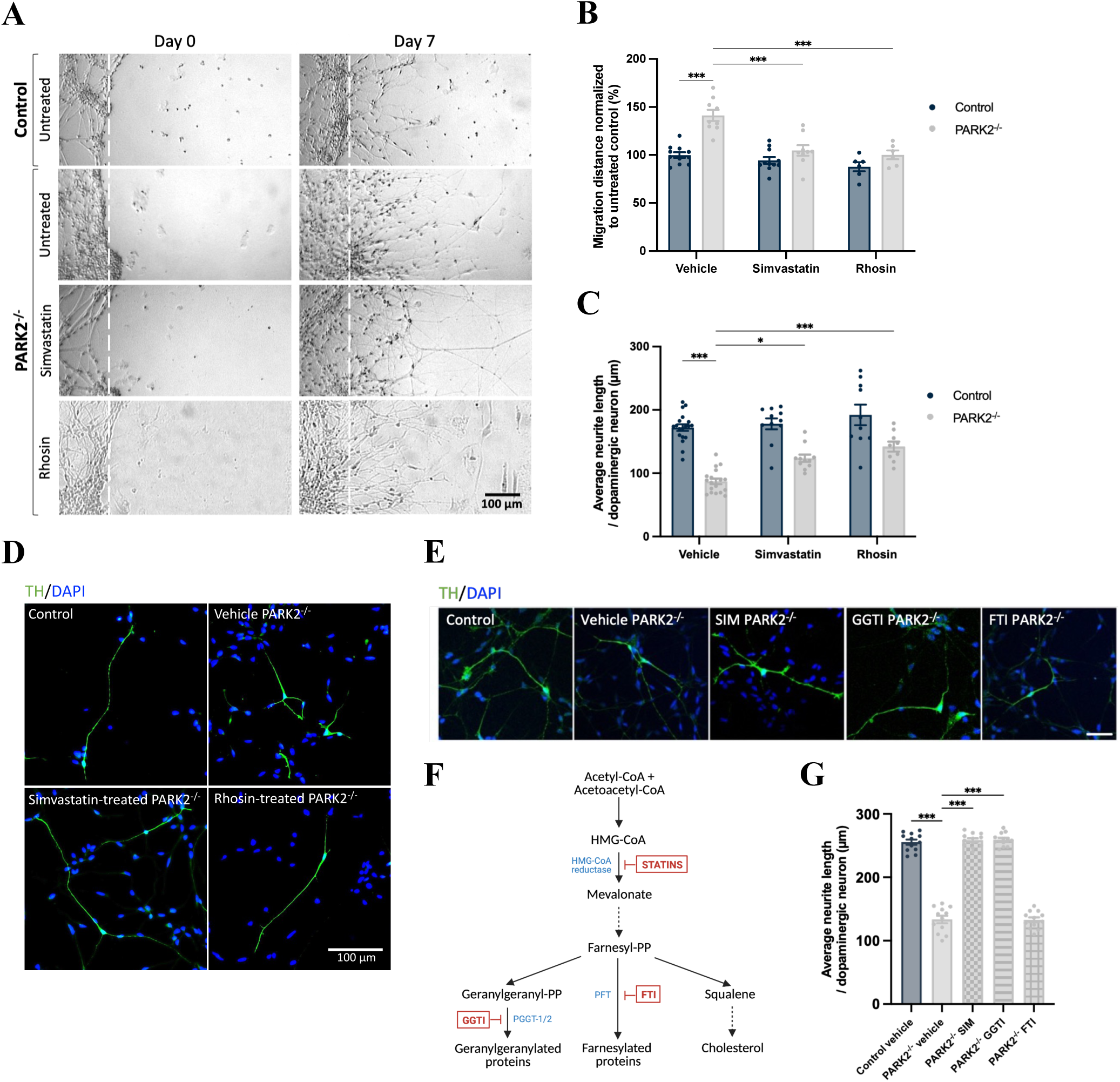
RhoA inhibition rescues increased migration of *PARK2*^-/-^ neurons and improve neurite outgrowth. A) Scratch assay was performed on day 18 (scratch day 0) of differentiation to evaluate migration of cell bodies into the scratch area after 7 days (scratch day 7). Scale bar = 100 µm. B) Quantification of the average migration distance revealed significantly increased migration of *PARK2*^-/-^ neurons, which was normalized by simvastatin and rhosin treatment. Mean ± SEM, n = 6-11 wells per group from 2 independent differentiations. Statistical analysis: two-way ANOVA, ***p < 0.001. C-D) Morphological analysis of TH^+^-dopaminergic neuronal precursors at day 13 of differentiation showed significantly reduced neurite length for *PARK2*^-/-^ cells, which were increased by simvastatin and rhosin treatment. Scale bar = 100 µm. Mean ± SEM, n = 9-19 coverslips per group from 3 independent differentiations. Statistical analysis: two-way ANOVA, *p < 0.05, ***p < 0.001. E-G) Inhibition of geranylgeranylation and farnesylation downstream of the HMG-CoA reductase in the mevalonate pathway using the geranylgeranyltransferase inhibitor (GGTI) and the farnesyltransferase inhibitor (FTI) showed that GGTI but not FTI phenocopied the effect of simvastatin on neurite outgrowth of *PARK2^-/-^* DA neurons. Scale bar = 50 µm. Mean ± SEM, n = 12 wells per group from 4 independent differentiations. Statistical analysis: one-way ANOVA, ***p < 0.001.

Neuritogenesis was investigated on TH^+^-DA precursors at day 13 of differentiation. *PARK2^-/-^* DA neurons displayed a reduced neurite length compared to isogenic controls, which could be increased by simvastatin treatment similar to the effect of direct RhoA inhibition after only three days of treatment (Fig. 2C-D). To further dissect the mechanism underlying simvastatin-mediated neurite outgrowth, we compared pharmacological inhibition of geranylgeranylation and farnesylation downstream in the mevalonate pathway (Fig. 2F). Treatment with the geranylgeranyltransferase inhibitor (GGTI) phenocopied the effect of simvastatin on neurite outgrowth of *PARK2^-/-^* DA neurons, whereas treatment with the farnesyltransferase inhibitor (FTI) had no effect (Fig. 2E and G). Both simvastatin and GGTI was confirmed to cause membrane dislocation of RhoA, while no effect was found for FTI (Fig. S2D-E). In summary, these data indicate that the effect of simvastatin treatment under our experimental conditions is indeed RhoA-mediated through reduced geranylgeranylation of RhoA, important for proper membrane localization and function of RhoA.

### RhoA inhibition rescues mitochondrial and lysosomal accumulation

We next sought to investigate the effect of simvastatin treatment on mitochondrial and lysosomal phenotypes in *PARK2^-/-^* neurons, as our recent studies found increased mitochondrial area and abnormal mitochondrial morphology and function associated with increased oxidative stress and autophagic-lysosomal perturbations upon parkin deficiency^27, 33, 34^.

Consistent with our previous findings, *PARK2^-/-^* neurons showed increased mitochondrial area evaluated by immunofluorescence staining for the OMM protein TOM20 (Fig. 3A-B). Both simvastatin treatment and direct RhoA inhibition were able to normalize the mitochondrial area to a level comparable to that of isogenic controls (Fig. 3A-B). However, treatment was not found to improve mitochondrial function as evaluated from ATP production (Fig. 3C).

**Figure 3.**
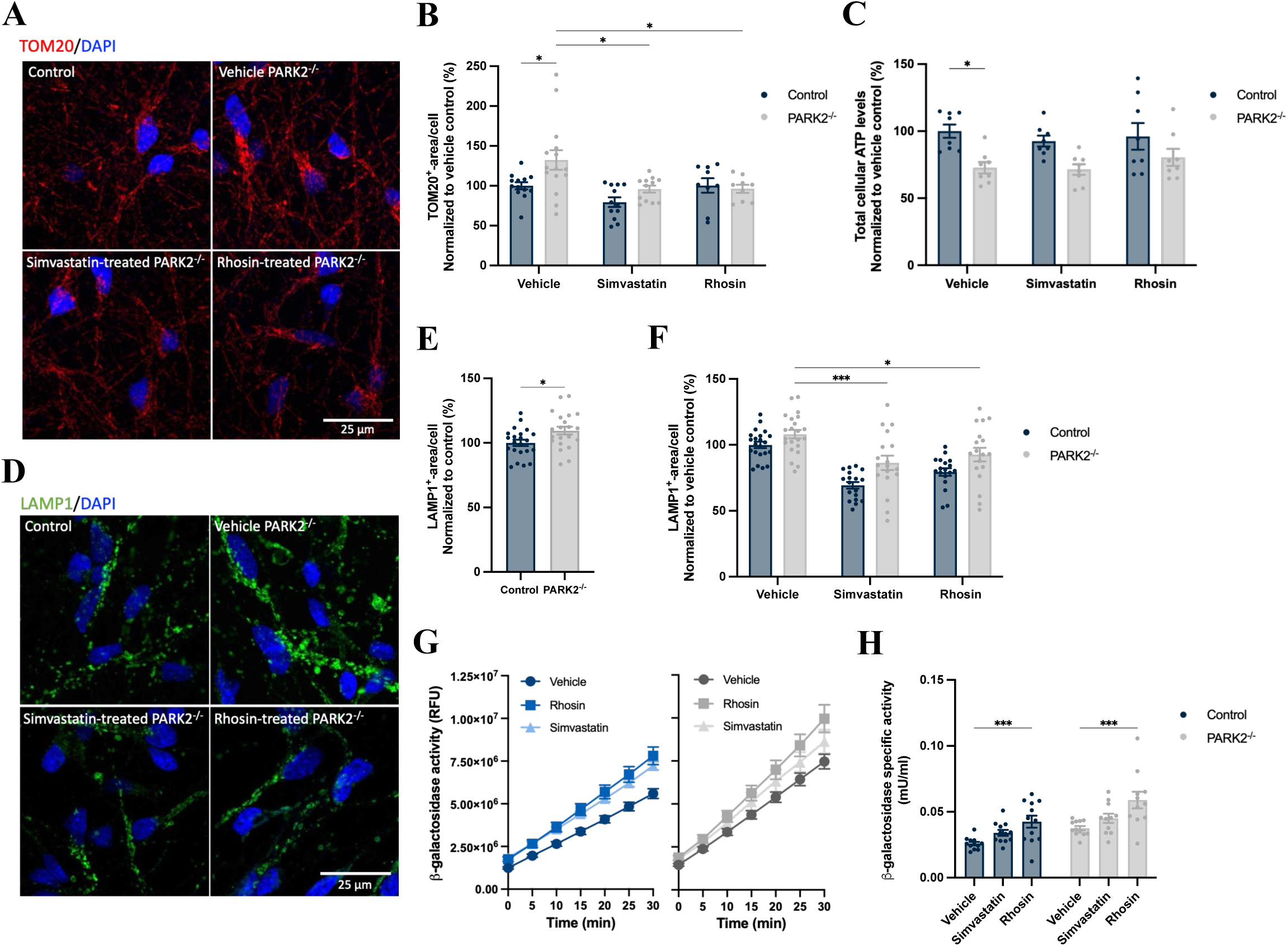
Simvastatin treatment and direct RhoA inhibition rescue increased mitochondrial and lysosomal area in *PARK2*^-/-^ neurons and increase lysosomal activity. A-B) Quantification of immunofluorescence staining for the outer mitochondrial membrane protein TOM20 (red) at day 35 of differentiation showed significantly increased TOM20^+^-mitochondrial area per cell in *PARK2*^-/-^ cultures relative to isogenic control cultures, which were normalized by statin and rhosin treatment. Mean ± SEM, n = 9-15 coverslips per group from 3-5 independent differentiations. Statistical analysis: two-way ANOVA, *p < 0.05. C) Total cellular ATP levels were significantly decreased in *PARK2*^-/-^ neurons relative to control but were not improved by simvastatin or rhosin treatment. Mean ± SEM, n = 8 wells per group from 2 independent differentiations. Statistical analysis: two-way ANOVA, *p < 0.05. D-E) Immunofluorescence staining for the lysosomal marker LAMP1 (green) and quantification showed significantly increased LAMP1^+^-lysosomal area in *PARK2*^-/-^ neurons relative to control. Mean ± SEM, n = 22 coverslips from 6 independent differentiations. Statistical analysis: Student’s t-test, *p < 0.05. F) Simvastatin and rhosin treatment significantly reduced the lysosomal area in *PARK2*^-/-^ neurons equal to the level in control neurons. Mean ± SEM, n = 15-22 coverslips per group from 6 independent differentiations. Statistical analysis: two-way ANOVA, *p < 0.05 ***p < 0.001. G) Activity of the lysosomal enzyme β-galactosidase was measured over time for control (blue) and *PARK2*^-/-^ (grey) with and without treatment. H) Calculation of specific β-galactosidase activity revealed increased activity upon direct RhoA inhibition and with a non-significant tendency for simvastatin. Mean ± SEM, n = 12 wells per group from 4 independent differentiations. Statistical analysis: one-way ANOVA, ***p < 0.001.

Immunofluorescence staining for the lysosomal marker LAMP1 revealed a slightly increased lysosomal area in *PARK2^-/-^* neurons (Fig. 3D-E), consistent with our previous findings^34^, and both simvastatin treatment and direct RhoA inhibition reduced the lysosomal area for both cell lines (Fig. 3D and F). Lysosomal function was evaluated by monitoring the activity of the lysosomal enzyme β-galactosidase (β-gal). β-gal activity was linear over the assay time, and a clear increase in activity was found for treated cells (Fig. 3H). Calculation of the specific β-gal activity confirmed increased activity upon treatment, but only significantly for direct RhoA inhibition. Together, these results suggest that RhoA inhibition rescues mitochondrial and lysosomal accumulation in *PARK2^-/-^*neurons, while only modestly improving lysosomal function and not restoring mitochondrial ATP production.

### RhoA inhibition changes mitophagy- and autophagy-related markers

To investigate if simvastatin and direct RhoA inhibition reduce mitochondrial area through modulation of mitophagy-related pathways, we investigated the expression levels of the ubiquitin-dependent mitophagy marker parkin and the ubiquitin-independent mitophagy marker NIX (BNIP3L) upon treatment. Both simvastatin treatment and direct RhoA inhibition increased parkin expression in control neurons (no expression found in *PARK2^-/-^* neurons, correlating with the gene knockout) (Fig. 4A and B). The same tendency was observed for NIX expression but was only significant for rhosin-treated control cultures (Fig. 4C and D). This suggests that RhoA inhibition may modulate pathways associated with both ubiquitin-dependent and ubiquitin-independent mitophagy^41^. We next investigated whether these changes were associated with alterations in general autophagy-related markers by analysing the expression levels of microtubule-associated protein 1 light chain 3B (LC3B) and P62. Direct RhoA inhibition markedly increased both total LC3B levels and LC3BII/I ratio, whereas simvastatin induced more modest changes (Fig. 4E-G). In parallel, P62 expression was elevated in *PARK2^-/-^* neurons compared with isogenic control neurons and was reduced following simvastatin treatment (Fig. 4H and I). Together, these findings indicate that RhoA inhibition is associated with changes in autophagy-related markers consistent with enhanced autophagic turnover.

**Figure 4.**
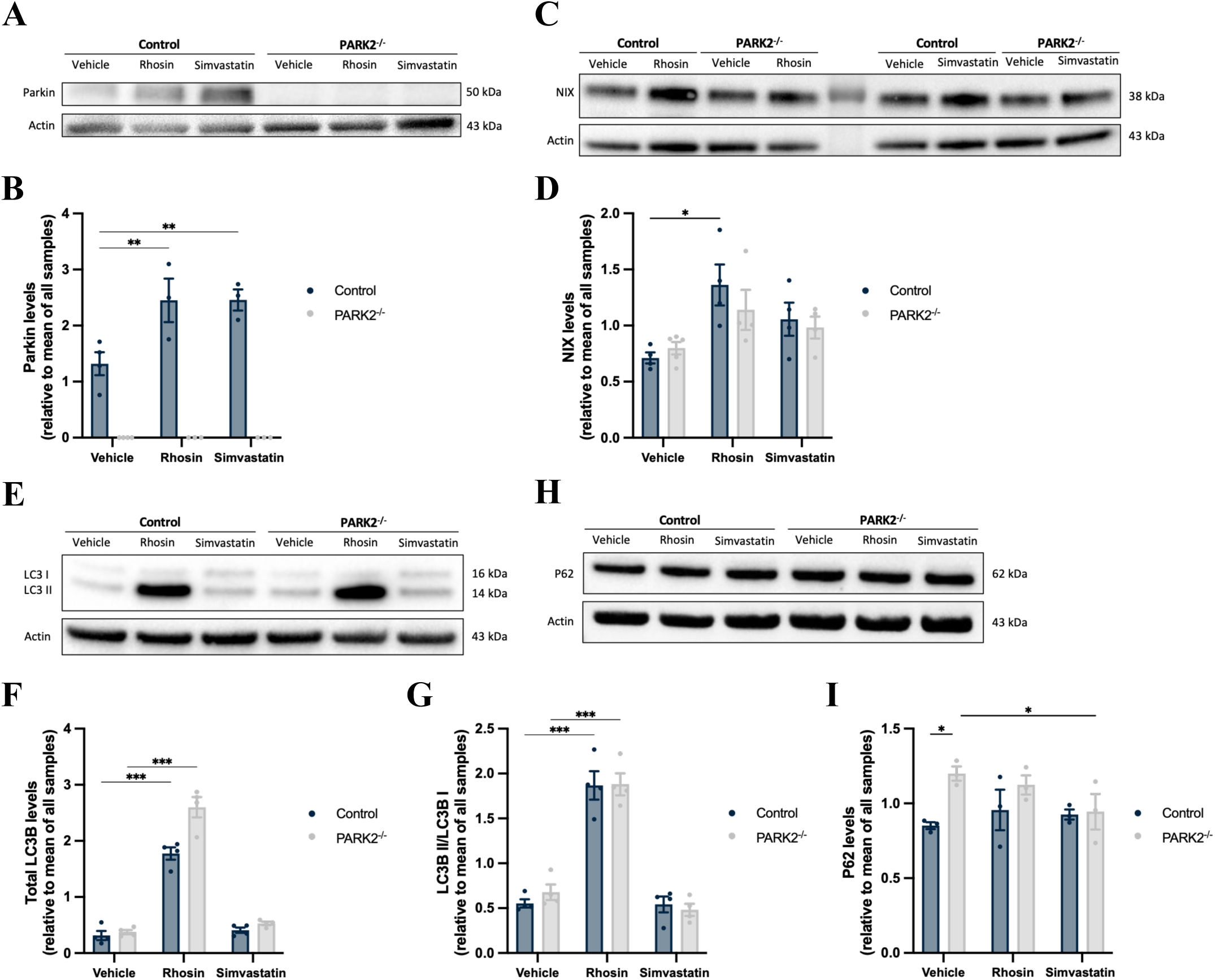
Treatment modulates mitophagy and autophagy. A-B) Western blotting for parkin showed significantly increased protein expression upon both simvastatin treatment and direct RhoA inhibition in control neurons and no parkin expression in *PARK2*^-/-^ neurons. C-D) Protein expression levels of NIX were increased in control cultures upon direct RhoA inhibition and showed a non-significant tendency for upregulation in the simvastatin-treated control group, as well as for treatment of *PARK2*^-/-^ neurons. E-G) Expression levels of LC3B quantified as both total LC3B levels (LC3B I + LC3B II) and LC3B turnover (LC3B II/ LC3B I) revealed significantly increased expression levels upon direct RhoA inhibition. H-I) P62 protein expression was elevated in *PARK2*^-/-^ neurons and reduced following simvastatin treatment. Protein expression levels were normalized to actin and are shown relative to the mean of all samples per differentiation. Mean ± SEM, n = 3-5 samples per group from 3-4 independent differentiations. Statistical analysis: two-way ANOVA, *p < 0.05, **p < 0.01, ***p < 0.001.

### RhoA inhibition reduces neuronal cell death and cytokine release

The effect of RhoA inhibition on cell death in the neuronal cultures was investigated for different treatment paradigms. Treatment was either initiated early (day 18) and cell death evaluated at day 45 of differentiation, or treatment was initiated late at day 38 and cell death evaluated at day 52 (Fig. 5A-D). At both timepoints, significantly increased cell death was observed in untreated *PARK2^-/-^*neuronal cultures evaluated by the release of lactate dehydrogenase (LDH) into the culture medium from necrotic cells (Fig. 5A and B). The increased LDH release was rescued by long-term direct RhoA inhibition, with shorter treatment showing a similar trend (Fig. 5A-B). Simvastatin treatment showed the same tendencies as direct RhoA inhibition, but its effect was not significant (Fig. 5A and B). Apoptotic cell death was evaluated by immunofluorescence staining for cleaved caspase 3 (casp3). A significantly increased number of casp3^+^-apoptotic cells was found in the *PARK2^-/-^* neuronal cultures compared to isogenic control cultures at both timepoints. In the early treatment paradigm, simvastatin significantly reduced the number of casp3+ cells, but not rhosin. In the late treatment paradigm, both simvastatin and direct RhoA inhibitior significantly reduced the number of casp3+ cells (Fig. 5C and D).

**Figure 5.**
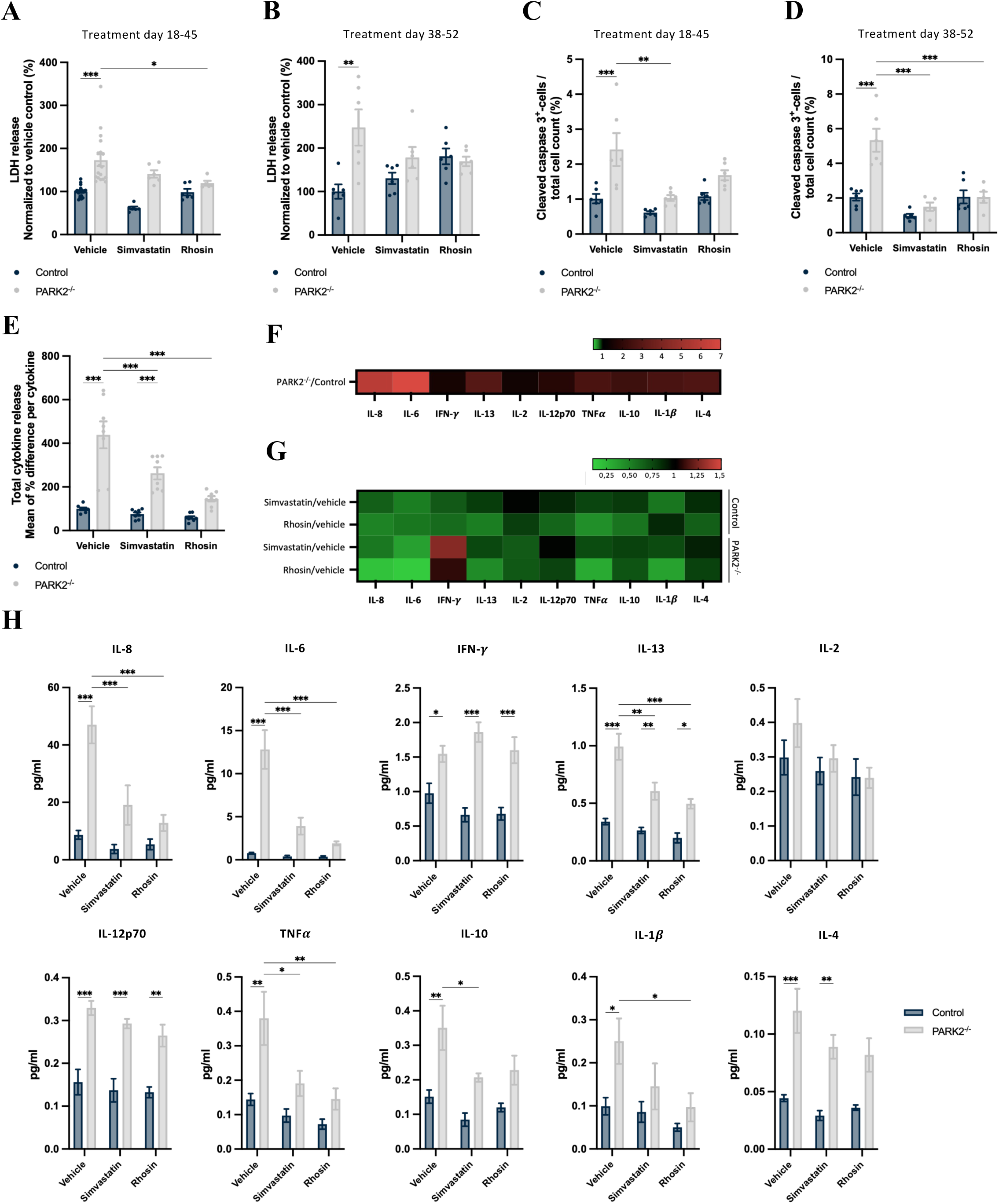
Treatment decreases cell death and cytokine release in *PARK2*^-/-^ cultures. A-B) Necrotic cell death was evaluated by LDH release and found increased in *PARK2*^-/-^ cultures compared to control cultures both at day 45 and 52. Direct RhoA inhibition decreased cell death in *PARK2*^-/-^ cultures. Mean ± SEM, n = 6-18 wells per group from 2-4 independent differentiations. C-D) Increased number of casp3^+^-apoptotic cells in *PARK2*^-/-^ neuronal cultures at day 45 and 52. Treatment from day 18-45 and 38-52 decreased apoptotic cell death in the *PARK2*^-/-^ cultures. Mean ± SEM, n = 5-6 from 2 independent differentiations. E) Total cytokine release, as evaluated by the mean fold change in cytokines levels per cytokine investigated relative to control levels, was increased in *PARK2*^-/-^ cultures and decreased by treatment. F-G) Heatmap showing relative levels of the individual cytokines investigated (IL-8, IL-6, IFN-γ, IL-13, IL-12p70, TNF-α, IL-10, IL-1β, and IL-4) in F) *PARK2*^-/-^ cultures relative to control and in G) treated groups relative to vehicle group for each cell line. H) Specific concentrations detected for the individual cytokines, showing increased levels for all cytokines except IL-2 in *PARK2*^-/-^ cultures and treatment-induced decrease in the levels of most cytokines. Mean ± SEM, n = 8 samples per group from 3 independent differentiations. Statistical analysis: two-way ANOVA, *p < 0.05, **p < 0.01, ***p < 0.001.

We next investigated if simvastatin treatment and direct RhoA inhibition could reduce modulate the levels of inflammatory cytokine secretion in the cultures. A marked increase in the total level of detected cytokines was found in *PARK2^-/-^* cultures compared to isogenic control, which was significantly decreased upon treatment (Fig. 5E-G). We observed that the release of interleukin (IL)-8, IL-6, interferon-γ (IFN-γ), IL-13, IL-12p70, tumour necrosis factor-α (TNF-α), IL-10, IL-1β, and IL-4 was significantly upregulated in *PARK2^-/-^* cultures (Fig. 5G and H). Both simvastatin treatment and direct RhoA inhibition decreased cytokine release for almost all cytokines detected, in both cell lines (Fig. 5G and H). The reduced release of IL-8, IL-6, IL-13, and TNF-α was significant for the *PARK2^-/-^* cultures upon both simvastatin treatment and direct RhoA inhibition. Interestingly, in contrast to the other cytokines assessed, no effect was found on the release of IFN-γ from *PARK2^-/-^*cultures was observed (Fig. 5H). Together, these findings demonstrate that RhoA inhibition reduces neuronal cell death and attenuates the pro-inflammatory cytokine profile in *PARK2^-/-^* cultures.

### Simvastatin treatment modulates disease-associated phenotypes in parkin PD patient-derived DA neurons

To understand the relevance of these findings from the *PARK2^-/-^* cell line, the effect of simvastatin treatment was further explored using parkin PD patient lines by evaluating a selection of the same phenotypic assays as performed for the isogenic *PARK2^-/-^* and control cultures.

iPSC lines from three compound heterozygous *PARK2* mutation carriers diagnosed with PD and five healthy control lines were differentiated into DA neuronal cultures with comparable efficiency (50%, Fig. S6A and C, cell line info in Table S1) using a modified dual-SMAD inhibition protocol as previously described^30^ (Fig. 6A). The midbrain identity of the DA neurons was confirmed by co-staining for FOXA2 and LMX1A (Fig. S6A, D and E). The maturity of the cultures was verified by a large expression of the mature neuronal marker MAP2 (90%) and expression of the pre- and postsynaptic markers synapsin and Homer 1, respectively (Fig. S6B and F).

**Figure 6.**
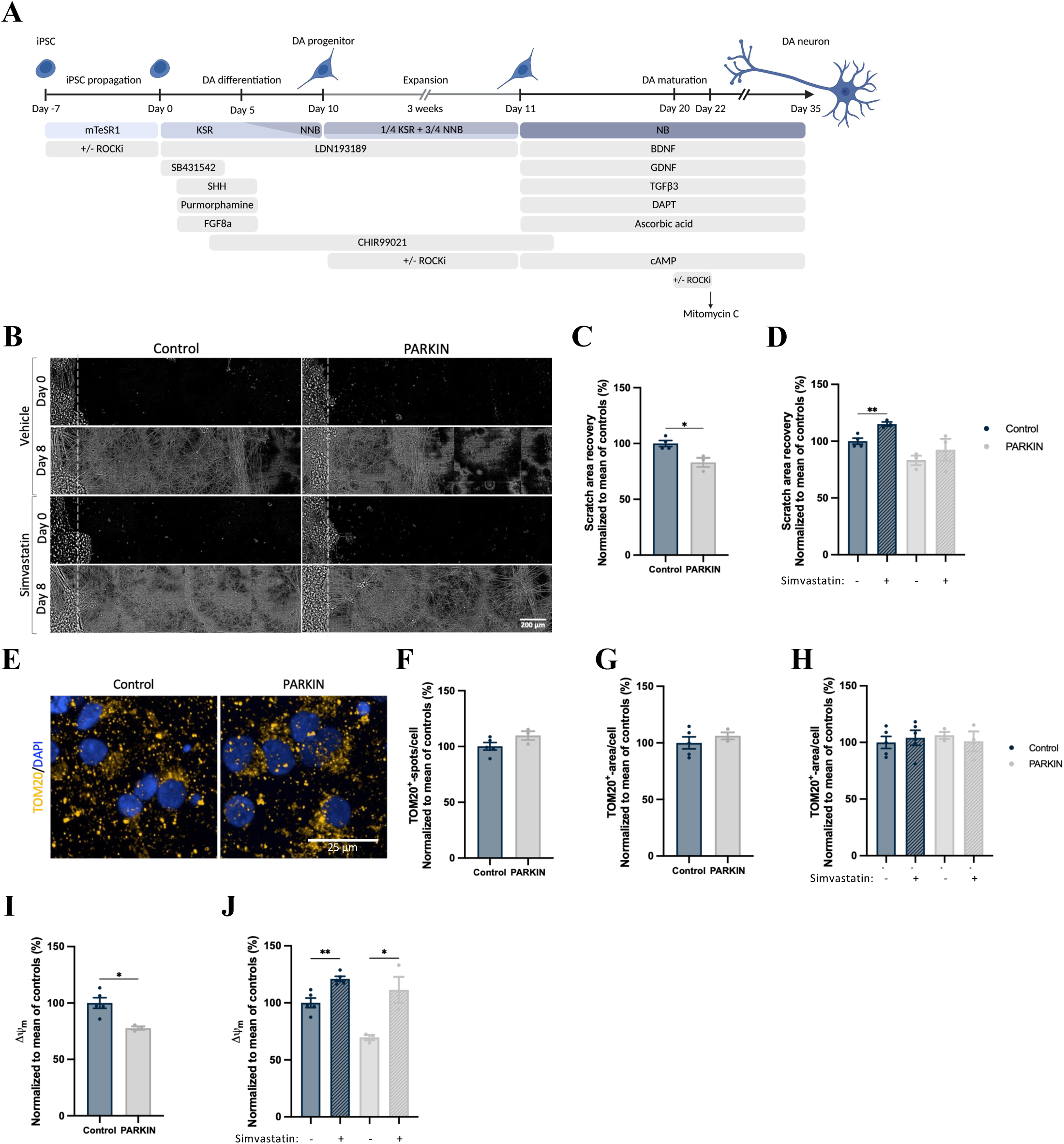
Effect of simvastatin treatment on parkin PD patient-derived DA cultures. A) Graphical overview of the dual-SMAD inhibition protocol applied for differentiating PD patient-derived iPSCs to dopaminergic neurons. B) Scratch assay was performed on differentiation day 30 (scratch day 0), and scratch area recovery was evaluated at differentiation day 38 (scratch day 8) as a measure of neurite outgrowth. C) Parkin PD patient-derived DA neurons showed decreased neurite outgrowth compared to healthy control neurons. Mean ± SEM, n = 4 control and 3 parkin cell lines, mean values from 3 independent differentiations. Statistical analysis: Student’s t-test, *p < 0.05. C) Simvastatin treatment significantly increased neurite outgrowth for control cultures, but not significantly for parkin cultures. Mean ± SEM, n = 4 control and 3 parkin cell lines, mean values from 3 independent differentiations. Statistical analysis: Student’s t-test comparing treatment group with untreated, **p < 0.01. E-H) Quantification of immunofluorescence staining for the outer mitochondrial membrane protein TOM20 (yellow) at day 35 of differentiation showed no difference in mitochondrial area or numbers in parkin DA neurons compared to healthy control neurons and with no effect of simvastatin treatment. Mean ± SEM, n = 5 control and 3 parkin cell lines, mean values from 3 independent differentiations. Statistical analysis: F-G) Student’s t-test, H) Student’s t-test comparing treatment group with untreated. I-J) Mitochondrial membrane potential (Δψm) was impaired for parkin PD patient DA neurons compared to healthy controls, and simvastatin treatment increased Δψm for both parkin and control DA neurons. Mean ± SEM, n = 5 control and 3 parkin cell lines, mean values from 2 independent differentiations. Statistical analysis: I) Student’s t-test, J) Student’s t-test comparing treatment group with untreated, *p < 0.05, **p < 0.01.

To assess neurite outgrowth, a scratch assay was performed on day 30 of differentiation, and scratch area recovery was evaluated at day 38 as a measure of neurite outgrowth (Fig. 6B). The assay revealed significantly reduced neurite outgrowth for the parkin PD patient DA cultures compared to healthy control cultures (Fig. 6B-C, individual cell line values are shown in Fig. S7A). Simvastatin treatment increased neurite outgrowth for control but not parkin cultures, although a non-significant tendency was observed (Fig. 6B and D, for dose response and toxicity see Fig. S8). To further investigate whether parkin loss of function causes neurite outgrowth defects, we knocked down (KD) *PARK2* using CRISPRi in the healthy control iPSC line WTC11 stably expressing the catalytically dead Cas9 (dCas9) system^28, 42^. Lentiviral single guide (sg) RNA transduction efficiency and differentiation characterization are found in the Supplementary Information (Fig. S9A-E). Neurite outgrowth was evaluated in sgRNA-expressing cells using a cytosolic FarRed reporter at day 2, 4, and 6 post-scratch (Fig. S9F-G). *PARK2* KD significantly reduced neurite outgrowth at day 6 post-scratch compared to non-targeting (NT) sgRNA transduced cells (Fig. S9G). Simvastatin treatment increased neurite outgrowth for *PARK2* KD neurons, evaluated at day 6 post-scratch (Fig. S9H). Interestingly, RhoA activity was also increased after *PARK2* KD (Fig. S9I).

We next investigated whether beneficial effects of simvastatin observed in the isogenic *PARK2^-/-^* model extended to mitochondrial phenotypes in patient-derived neurons. While TOM20 staining revealed no difference in mitochondrial number or area between parkin PD patient and control cultures, indicating preserved mitochondrial content (Fig. 6E-H and S7B-C), mitochondrial membrane potential (Δψm) was significantly reduced in parkin PD patient-derived neurons compared with healthy controls (Fig. 6I and J, and S7D). Importantly, simvastatin significantly increased Δψm in both control and parkin PD patient-derived neurons, indicating improved mitochondrial function despite the absence of detectable changes in mitochondrial abundance.

In summary, these findings demonstrate that simvastatin confers beneficial effects in both isogenic and patient-derived *PARK2* models. While the magnitude of the response varied among patient-derived lines, likely reflecting differences in *PARK2* genotype (including compound heterozygous mutations) and genetic background, the overall direction of the effects was consistent across models and supports the translational relevance of RhoA inhibition in parkin-associated PD.

### Increased RhoA activity in PD patient-derived DA neurons

Given the observed perturbations in RhoA in isogenic *PARK2^-/-^* iPSC-derived neurons, we sought to understand if RhoA activity is altered across genetic PD lines. We performed a RhoA activity screen of PD patient iPSC-derived DA neurons with different PD-related gene mutations (PINK1 loss of function, parkin loss of function, *LRRK2* (G2019S), *LRRK2* (R1441C), *GBA* (L44P), *GBA* (N370S), *SNCA* (A53T), and *SNCA* (triplication)) (Cell line info in Table S1). A total of 32 PD patient and control iPSC lines were differentiated to DA neurons for 35 days using a dual-SMAD inhibition protocol (Fig. 6A). RhoA activity was found upregulated across many of the PD patient lines relative to healthy control cell lines, but not all (Fig. 7A). Consistent with our observations in isogenic *PARK2*^-/-^ lines, we observed a significant upregulation in neuronal cultures from PD patients with PARK2 mutations (Fig. 7B). Furthermore, we also observed increased RhoA activity in neurons from PD patients with *PINK1* mutations and patients with a *SNCA* A53T mutation relative to healthy controls. In contrast, patients with *LRRK2* (G2019S), *LRRK2* (R1441C), *GBA* (L44P), *GBA* (N370S), and *SNCA* (triplication) mutations did not show significantly increased RhoA activity relative to healthy controls. These findings demonstrate that increased RhoA activity is present across multiple genetic forms of PD, supporting dysregulated RhoA signalling as a common pathogenic mechanism and potential disease-modifying therapeutic target.

**Figure 7.**
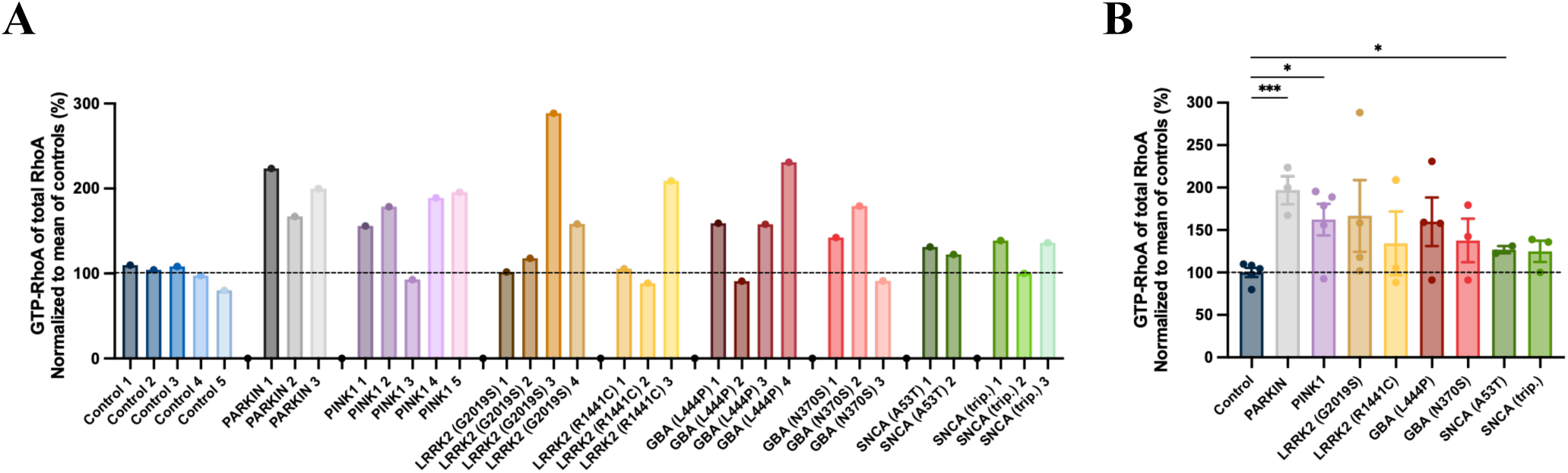
Increased RhoA activity in PD patient-derived DA neurons. A) 32 different PD patient and control iPSC-derived DA neuronal lines were screened for RhoA activity levels. B) Mean RhoA activity levels across the different PD-related gene mutations (PINK1 loss of function, parkin loss of function, *LRRK2* (G2019S), *LRRK2* (R1441C), *GBA* (L44P), *GBA* (N370S), *SNCA* (A53T), and *SNCA* (triplication)) show increased levels across all lines relative to control. Mean ± SEM, n = 3-5 cell lines per group. Statistical analysis: one-way ANOVA, *p < 0.05, ***p < 0.001.

## DISCUSSION

Statins are of increasing interest for their potential therapeutic effect in the treatment of PD^21, 43, 44^. Previous epidemiological studies provide inconclusive data, however, with some suggesting that statins reduce the risk of developing PD^45, 46^ and others reporting no such association^47–49^ or even an increased risk^50, 51^. Clinical studies also show conflicting results, reflecting heterogeneity in clinical trial outcomes^52^. A randomized clinical trial of simvastatin as a disease-modifying treatment for patients with moderate PD found no significant clinical benefit (PD STAT)^53^, whereas a clinical trial of lovastatin in early-stage PD showed a tendency to reduce the motor symptom progression^54^. These conflicting results likely reflect differences in patient stratification, disease stage, lipophilic vs. hydrophilic statins, and duration effects, highlighting the need for further in-depth mechanistic studies.

Previous *in vitro* and *in vivo* studies have shown that statins do exert neuroprotective effects in addition to their cholesterol-lowering effect^55–58^, by reducing the levels of isoprenoid intermediates in the cholesterol biosynthetic pathway, which is important for proper membrane localization and function of the Rho GTPases including RhoA^19^. Lipophilic statins can cross the blood-brain-barrier, and simvastatin reaches peak concentrations of 300-500 nM in mice brains^24^. Simvastatin has been shown to inhibit RhoA prenylation and membrane association at clinically relevant doses of simvastatin as low as 50 nM^19, 59^. While several *in vitro* studies have reported isoprenoid-dependent effects of statins in neurons and glial cells using high statin concentrations (usually between 1-10 µM but up to 100 µM)^60–62^, the effect of statins at lower and more clinically relevant doses is not well studied.

In the present study, we show that simvastatin at a clinically relevant dose of 300 nM modulates several important processes involved in driving neurodegeneration in PD. We applied human iPSC-derived DA neurons both PD patient-derived and isogenic *PARK2^-/-^* cell lines and included direct RhoA inhibition using rhosin as a positive control. Consistent with our previous findings, *PARK2^-/-^* neurons display increased RhoA activity compared to isogenic control neurons^18^. Strikingly, by performing the first systematic screen of RhoA activity across 32 iPSC-derived DA neuron lines representing multiple genetic forms of PD (PINK1 loss of function, parkin loss of function, *LRRK2* (G2019S), *LRRK2* (R1441C), *GBA* (L44P), *GBA* (N370S), *SNCA* (A53T), and *SNCA* (triplication)) and healthy controls, we show for the first time that RhoA activity was increased across many of the PD patient lines compared to healthy control lines. This suggests that targeting RhoA signalling might be a relevant target in genetic forms of PD. While increased RhoA activity has previously only been reported in neurotoxin-induced mouse and rat models of PD^10, 13, 14, 63^ and our own report on increased RhoA activity in human *PARK2^-/-^* neurons^18^, this is the first RhoA activity screen reported on human iPSC-derived DA neurons from patients with PD-related mutations. To further validate RhoA signalling as a target for PD in general, future studies need to address whether increased RhoA activity is also found in sporadic PD. As not all PD patient-derived cell lines in our screen showed increased RhoA activity (even within the same mutation group), it might be necessary to stratify patients based on RhoA activity levels to determine eligibility for a RhoA targeting treatment such as simvastatin. Major functions of RhoA signalling are to regulate actin cytoskeleton dynamics and RhoA activation, promote migration of neuronal precursor cells, and cause axonal and dendritic retraction and spine and synapse loss^6, 8^. In accordance with this, our previously reported pathway analysis of proteomic and PTM-omic changes identified for *PARK2^-/-^* vs. isogenic control neurons predicted changes in cell migration and neurite outgrowth. These were confirmed by *in vitro* and *in vivo* assays and morphological analyses of *PARK2^-/-^* DA neurons and could be rescued by RhoA inhibition^18^. In the current study, we show that treatment with 300 nM simvastatin reduces migration and increases neurite outgrowth of *PARK2^-/-^*neurons equal to the effect of direct RhoA inhibition using rhosin. Similarly, parkin PD patient-derived DA neurons showed decreased neurite outgrowth compared to healthy control DA neurons.

Our findings of neurite outgrowth-enhancing effects of simvastatin treatment are consistent with previous studies that applied higher simvastatin concentrations of 1-10 µM^56, 64^. Mechanistic studies investigating products of the mevalonate pathway that mediate the effect of statins on neurite outgrowth have shown that addition of mevalonate, FPP, and GGPP but not cholesterol prevented the effects of statins on neurite outgrowth and membrane association of RhoA^62, 64^. Moreover, inhibition of the geranylgeranyltransferase but not farnesyltransferase mimicked the effect of statins^62^. In accordance with these observations, the simvastatin concentration used in the present study did not reduce intracellular cholesterol levels in the cell cultures, and treatment with the fibrate gemfibrozil (which does not prevent RhoA prenylation when lowering cholesterol levels) did not reduce migration of *PARK2^-/-^*neurons. Taken together, this strongly suggests that the statin-mediated induction of neurite outgrowth relies on the inhibition of RhoA prenylation. This is supported by our finding that simvastatin reduced membrane-associated RhoA levels similar to GGTI, whereas FTI treatment had no effect, in line with a requirement for geranylgeranylation in RhoA membrane association. These findings are also consistent with studies showing that statins can directly influence membrane biophysical properties, including lipid packing in phospholipid bilayers^65, 66^, which may also affect membrane association and localization of signalling proteins such as RhoA.

Neurite outgrowth defects have previously also been reported in iPSC-derived neurons from *LRRK2*, *GBA,* and sporadic PD patients^31, 67, 68^. Synaptic and axonal degeneration is one of the earliest pathological events in PD^69^, so a modulatory effect of RhoA inhibition on neurite outgrowth suggests that targeting RhoA signalling could have potential in early disease intervention in PD. Whether this is indicative of statin treatment being mostly effective for early-stage PD needs further validation in preclinical models, but it highlights a potential need for refining treatment initiation and duration in clinical studies.

Neuroinflammation mediated by the release of pro-inflammatory cytokines and oxidative stress is a prime contributor to the pathogenesis of PD^5, 70^. Notably, simvastatin treatment was found to significantly decrease the elevated levels of almost all cytokines detected in the *PARK2^-/-^* cultures. The increased cytokine release in the *PARK2^-/-^* cultures is most likely from the astrocyte population in the cultures rather than from the neurons. Neuroinflammation is driven by the activation of glial cells, mainly microglia but also astrocytes, and of peripheral infiltrating T lymphocytes^71^. Our differentiated cultures do not contain microglial cells as they have a mesodermal origin^72^, while the iPSC-derived NSCs are ectoderm-derived. Although the proportion of astrocytes was similar between *PARK2^-/-^* and control cultures, we qualitatively observed more pronounced GFAP staining and hypertrophic-like astrocytic morphology in *PARK2^-/-^* cultures. In line with this, we have previously reported increased protein expression of GFAP and vimentin in *PARK2^-/-^* cultures ^18^, supporting a potential increase in astrocytic reactivity. As reactive astrocytes are known to secrete proinflammatory cytokines^35^, this may contribute to the elevated cytokine levels observed in the *PARK2^-/-^*cultures. RhoA signalling is highly involved in determining astrocyte morphology and activation state^73^, and inhibition of RhoA signalling decreases reactive gliosis^74^. Similarly, simvastatin treatment has been shown to decrease astrocyte reactivity and cytokine release in neurotoxin-induced PD models^75, 76^.

Although neuroinflammation is believed to be a key player in the progression of PD, it is most likely not the initial trigger^77, 78^. Increasing evidence points towards mitochondrial dysfunction as a key determinant of DA neuronal susceptibility in both sporadic and familial PD^79^. Our recent studies of *PARK2^-/-^* neurons have shown increased mitochondrial area, abnormal morphology, and impaired mitochondrial membrane potential associated with decreased ATP production and metabolomic disturbances^27, 33^. In the current study, we show that simvastatin treatment and direct RhoA inhibition decrease the mitochondrial area in *PARK2^-/-^* neurons to a level comparable with that of isogenic control neurons, but treatment did not increase mitochondrial function as evaluated by ATP production. In parkin PD patient-derived neuronal cultures, we found no differences in either mitochondrial area or numbers, but the mitochondrial membrane potential was significantly decreased in the parkin PD patient neurons compared to healthy controls. Simvastatin treatment significantly increased the mitochondrial membrane potential for both controls and parkin patient cultures. To our knowledge, this is the first report of statin-mediated improvements of mitochondrial membrane potential deficits in a PD model, but similar findings have been reported for other neurodegenerative diseases. Both simvastatin treatment and fasudil-mediated inhibition of the RhoA downstream effector ROCK improved mitochondrial membrane potential deficits and protected against striatal neurodegeneration in the 3-nitropropionic acid-induced Huntington’s disease rat model^80^. Similarly, lovastatin treatment increased mitochondrial membrane potential and protected oligodendrocytes against cytokine toxicity in an *in vitro* multiple sclerosis model^81^. It has been proposed that the underlying mechanism linking the anti-oxidative, anti-inflammatory, and mitochondrial-improving effects of statin treatment is through PPARα-dependent transcriptional activation^82^, and it has been shown that statin-mediated inhibition of RhoA/ROCK signalling induces PPARα activation^81^.

Mitochondrial homeostasis is highly linked to proper lysosomal function^83–86^, and we have previously established lysosomal alterations in *PARK2^-/-^* neurons^34^. In the present study, we found that simvastatin treatment and direct RhoA inhibition reversed the increased lysosomal area in *PARK2^-/-^* neurons and improved lysosomal function as evaluated by β-gal activity. Accordingly, simvastatin treatment has previously been reported to improve lysosome function via enhancing lysosome biogenesis and its autophagic turnover in endothelial cells^87^. Our findings of simvastatin treatment and RhoA inhibition normalizing the mitochondrial and lysosomal areas in *PARK2^-/-^* neurons might therefore suggest modulation of mitophagy- and autophagy-related pathways. However, as autophagic and mitophagic flux were not directly assessed (e.g., using mt-Keima or lysosomal inhibition approaches), future studies are needed to determine whether RhoA inhibition enhances functional mitophagy. It was recently shown that ROCK inhibitors promote parkin-mediated mitophagy by Akt-mediated activation of hexokinase 2 (HK2), a positive regulator of parkin^88^. Indeed, we found upregulation of parkin in controls cells upon RhoA inhibition and simvastatin treatment, but NIX levels were also increased by treatment in *PARK2^-/-^* and controls neurons, consistent with activation of signalling pathways associated with both ubiquitin-dependent and ubiquitin-independent mitophagy^89^.

Likewise, direct inhibition of RhoA signalling altered autophagy-related markers, as evidenced by increased total LC3B levels and LC3B-II/I ratios together with reduced P62 accumulation, findings that are consistent with enhanced autophagic turnover and increased autophagosome formation. Although simvastatin induced more modest changes in LC3B compared with direct RhoA inhibition, the reduction in P62 levels together with normalization of mitochondrial and lysosomal phenotypes suggests that simvastatin may also improve autophagic and mitophagic turnover. Previous studies have similarly reported increased autophagy following statin treatment at higher concentrations^87, 90^. In addition, it cannot be ruled out that simvastatin may affect lysosomal and metabolic signalling pathways upstream of autophagy, including mTOR–TFEB signalling and sterol-sensing pathways such as the NPC1/mTOR axis, thereby influencing transcriptional regulation of lysosomal and inflammatory genes and broader sterol homeostasis^87, 91–93^.

In summary, neurite outgrowth defects could be related to mitochondrial dysfunctions^94^ that cause lack in energy production, which the highly metabolomic active DA neurons are particularly vulnerable towards^95^. Therefore, the neuroprotective mechanism of simvastatin treatment might be linked to its mitophagy-enhancing effects. Moreover, autophagosome biogenesis and trafficking are also highly dependent on cytoskeletal dynamics^96^ that are regulated by RhoA signalling. Through ROCK, RhoA signalling is reported to regulate intracellular redistribution of lysosomes^97^, suggesting that RhoA signalling might regulate trafficking from early autophagosomes to late autolysosomes important for the autophagy process, which perhaps explains the observed effect of simvastatin treatment. The exact mechanism needs further investigation, but as multiple cellular processes have been implicated in the pathogenesis of PD, it is highly interesting that we find a positive effect of simvastatin treatment and direct RhoA inhibition on several disease-relevant cellular phenotypes.

## CONCLUSION

We believe this is the most comprehensive *in vitro* study investigating the disease-modifying effects of simvastatin treatment for PD. By applying human iPSC-derived neurons with *PARK2^-/-^* and isogenic control neurons, we show that treatment with simvastatin at a physiologically relevant dose, including direct RhoA inhibition as a positive control, shows neuroprotective effects through several PD-relevant mechanisms. These include DA neurite outgrowth, anti-inflammatory action, mitochondrial and lysosomal phenotypes, modulation of mitophagy- and autophagy-related markers, and ultimately decreased cell death. Moreover, we provide the very first RhoA activity screen of PD patient iPSC-derived DA neurons with different PD-related gene mutations (PINK1 loss of function, parkin loss of function, *LRRK2* (G2019S), *LRRK2* (R1441C), *GBA* (L44P), *GBA* (N370S), *SNCA*

(A53T), and *SNCA* triplication) and healthy controls, and we show that RhoA activity was increased for many of the PD patient lines compared to healthy control lines, though not all. These data strongly suggest that targeting RhoA signalling, e.g. by simvastatin treatment, might be a potential disease-modifying strategy for PD, but it also highlights the need for patient stratification based on RhoA activity levels prior to clinical trial recruitment.

## LIST OF ABBREVIATIONS

ANOVA: Analysis of variance
ATP: Adenosine triphosphate
BDNF: Brain-derived neurotrophic factor
BCA: Bicinchoninic acid
CCCP: Carbonyl cyanide 3-chlorophenylhydrazone
CHIR99021: GSK-3β inhibitor (small molecule)
CRISPRi: Clustered regularly interspaced short palindromic repeats interference
DA: Dopaminergic
DAPI: 4′,6-diamidino-2-phenylindole dihydrochloride
DMSO: Dimethyl sulfoxide
DOPA: L-3,4-dihydroxyphenylalanine
FPP: Farnesyl pyrophosphate
FTI: Farnesyl transferase inhibitor
GBA: Glucocerebrosidase
GDNF: Glial cell line-derived neurotrophic factor
GGTI: Geranylgeranyl transferase inhibitor
GFAP: Glial fibrillary acidic protein
HMG-CoA: 3-Hydroxy-3-methylglutaryl-coenzyme
A IF: Immunofluorescence
IFN-γ: Interferon gamma
IL: Interleukin
iPSC: Induced pluripotent stem cell
KD: Knockdown
LC3B: Microtubule-associated protein 1 light chain 3B
LDH: Lactate dehydrogenase
LRRK2: Leucine-rich repeat serine/threonine-protein kinase 2
MAP2: Microtubule-associated protein 2
MITO: Mitochondrial (context-specific abbreviation used in assays)
MPTP: 1-Methyl-4-phenyl-1,2,3,6-tetrahydropyridine
NEAA: Non-essential amino acids
NIX: BCL2/adenovirus E1B 19 kDa protein-interacting protein 3-like
NNB: Neurobasal-based medium (context-specific)
NT: Non-targeting (sgRNA control)
OMM: Outer mitochondrial membrane PBS Phosphate-buffered saline
PD: Parkinson’s disease
PFA: Paraformaldehyde
PINK1: PTEN-induced putative kinase 1
PLo: Poly-L-ornithine
PVDF: Polyvinylidene difluoride
PTM: Post-translational modification
RhoA: Ras homolog gene family member A
ROS: Reactive oxygen species
RT: Room temperature
SB431542: TGF-β receptor inhibitor
SHH: Sonic hedgehog
SNCA: α-synuclein gene
SNpc: Substantia nigra pars compacta
TH: Tyrosine hydroxylase
TNF-α: Tumor necrosis factor alpha
TBS: Tris-buffered saline
TGFβ3: Transforming growth factor beta 3
VDAC: Voltage-dependent anion channel (context-specific, if used)
V-PLEX: MesoScale multiplex cytokine assay platform
WTC11: Wild-type control iPSC line 11
Y27632: ROCK inhibitor (small molecule)

## DECLARATIONS

### ETHICS APPROVAL

All use of human stem cells was performed in accordance with the Danish national regulations, the International Society for Stem Cell Research (ISSCR) and the National Health Service, Health Research Authority, NRES Committee South Central, Berkshire, UK, REC 10/H0505/71.

### CONSENT FOR PUBLICATION

Not applicable.

### AVAILABILITY OF DATA AND MATERIALS

The dataset used and/or analysed during the current study are available from the corresponding author on request.

### AUTHORS’ CONTRIBUTIONS

S.I.S., M.R., J.O. B.R., and M.M. designed the research; S.I.S., M.R., I.L.K.S., N.F.B.J., E.B.C., M. K. and J.O. performed experiments; A.D., D.W., R.W.M. contributed new reagents or analytic tools; S.I.S., M.R., I.L.K.S., N.F.B.J., E.B.C., J.O. and L.S.W. analysed data, S.I.S, J.O. and M.M. wrote the manuscript; K.F., B.R., and M.B. revised the manuscript. All authors read and approved the final manuscript.

## ACKNOWLEDGMENTS

The authors would like to thank Nadine Becker-von Buch for excellent technical assistance and Dr. Claire Gudex for proofreading the manuscript.

## REFERENCES

1. Kalia, L.V. & Lang, A.E. Parkinson’s disease. Lancet 386, 896–912 (2015).

2. Balestrino, R. & Schapira, A.H.V. Parkinson disease. Eur J Neurol 27, 27–42 (2020).

3. Corti, O., Lesage, S. & Brice, A. What genetics tells us about the causes and mechanisms of Parkinson’s disease. Physiol Rev 91, 1161–1218 (2011).

4. Lesage, S. & Brice, A. Parkinson’s disease: from monogenic forms to genetic susceptibility factors. Hum Mol Genet 18, R48–59 (2009).

5. Poewe, W. et al. Parkinson disease. Nat Rev Dis Primers 3, 17013 (2017).

6. Schmidt, S.I., Blaabjerg, M., Freude, K. & Meyer, M. RhoA Signaling in Neurodegenerative Diseases. Cells 11 (2022).

7. Ravichandran, N. et al. New insights on the regulators and inhibitors of RhoA-ROCK signalling in Parkinson’s disease. Metab Brain Dis 40, 90 (2025).

8. Stankiewicz, T.R. & Linseman, D.A. Rho family GTPases: key players in neuronal development, neuronal survival, and neurodegeneration. Front Cell Neurosci 8, 314 (2014).

9. Arrazola Sastre, A., et al. Small GTPases of the Ras and Rho Families Switch on/off Signaling Pathways in Neurodegenerative Diseases. Int J Mol Sci 21 (2020).

10. Villar-Cheda, B. et al. Involvement of microglial RhoA/Rho-kinase pathway activation in the dopaminergic neuron death. Role of angiotensin via angiotensin type 1 receptors. Neurobiol Dis 47, 268–279 (2012).

11. Tonges, L. et al. Inhibition of rho kinase enhances survival of dopaminergic neurons and attenuates axonal loss in a mouse model of Parkinson’s disease. Brain 135, 3355–3370 (2012).

12. Borrajo, A., Rodriguez-Perez, A.I., Villar-Cheda, B., Guerra, M.J. & Labandeira-Garcia, J.L. Inhibition of the microglial response is essential for the neuroprotective effects of Rho-kinase inhibitors on MPTP-induced dopaminergic cell death. Neuropharmacology 85, 1–8 (2014).

13. Sanchez, M., Gastaldi, L., Remedi, M., Caceres, A. & Landa, C. Rotenone-induced toxicity is mediated by Rho-GTPases in hippocampal neurons. Toxicol Sci 104, 352–361 (2008).

14. Niu, M., Xu, R., Wang, J., Hou, B. & Xie, A. MiR-133b ameliorates axon degeneration induced by MPP(+) via targeting RhoA. Neuroscience 325, 39–49 (2016).

15. Mattii, L. et al. Rho-inhibition and neuroprotective effect on rotenone-treated dopaminergic neurons in vitro. Neurotoxicology 72, 51–60 (2019).

16. Bloem, B.R., Okun, M.S. & Klein, C. Parkinson’s disease. Lancet 397, 2284–2303 (2021).

17. Malpartida, A.B., Williamson, M., Narendra, D.P., Wade-Martins, R. & Ryan, B.J. Mitochondrial Dysfunction and Mitophagy in Parkinson’s Disease: From Mechanism to Therapy. Trends Biochem Sci 46, 329–343 (2021).

18. Bogetofte, H. et al. Perturbations in RhoA signalling cause altered migration and impaired neuritogenesis in human iPSC-derived neural cells with PARK2 mutation. Neurobiol Dis 132, 104581 (2019).

19. Ostrowski, S.M. et al. Simvastatin inhibits protein isoprenylation in the brain. Neuroscience 329, 264–274 (2016).

20. Al-Kuraishy, H.M. et al. The effects of cholesterol and statins on Parkinson’s neuropathology: a narrative review. Inflammopharmacology 32, 917–925 (2024).

21. Fracassi, A. et al. Statins and the Brain: More than Lipid Lowering Agents? Curr Neuropharmacol 17, 59–83 (2019).

22. Reddy, J.M., Raut, N.G.R., Seifert, J.L. & Hynds, D.L. Regulation of Small GTPase Prenylation in the Nervous System. Mol Neurobiol 57, 2220–2231 (2020).

23. Ye, Q. et al. Role of Rho-associated kinases and their inhibitor fasudil in neurodegenerative diseases. Front Neurosci 18, 1481983 (2024).

24. Johnson-Anuna, L.N. et al. Chronic administration of statins alters multiple gene expression patterns in mouse cerebral cortex. J Pharmacol Exp Ther 312, 786–793 (2005).

25. Shaltouki, A. et al. Mitochondrial alterations by PARKIN in dopaminergic neurons using PARK2 patient-specific and PARK2 knockout isogenic iPSC lines. Stem Cell Reports 4, 847–859 (2015).

26. Fernandes, H.J. et al. ER Stress and Autophagic Perturbations Lead to Elevated Extracellular alpha-Synuclein in GBA-N370S Parkinson’s iPSC-Derived Dopamine Neurons. Stem Cell Reports 6, 342–356 (2016).

27. Okarmus, J. et al. Identification of bioactive metabolites in human iPSC-derived dopaminergic neurons with PARK2 mutation: Altered mitochondrial and energy metabolism. Stem Cell Reports 16, 1510–1526 (2021).

28. Williamson, M. G., et al. USP30 inhibition improves mitochondrial health through both PINK1-dependent and independent mechanisms. bioRxiv (2025). doi:10.1101/2025.02.03.636341.

29. Connor-Robson, N. et al. An integrated transcriptomics and proteomics analysis reveals functional endocytic dysregulation caused by mutations in LRRK2. Neurobiol Dis 127, 512–526 (2019).

30. Williamson, M.G. et al. Mitochondrial dysfunction and mitophagy defects in LRRK2-R1441C Parkinson’s disease models. Hum Mol Genet (2023).

31. Bogetofte, H. et al. Post-translational proteomics platform identifies neurite outgrowth impairments in Parkinson’s disease GBA-N370S dopamine neurons. Cell Rep 42, 112180 (2023).

32. Zambon, F. et al. Cellular alpha-synuclein pathology is associated with bioenergetic dysfunction in Parkinson’s iPSC-derived dopamine neurons. Hum Mol Genet 28, 2001–2013 (2019).

33. Bogetofte, H. et al. PARK2 Mutation Causes Metabolic Disturbances and Impaired Survival of Human iPSC-Derived Neurons. Front Cell Neurosci 13, 297 (2019).

34. Okarmus, J. et al. Lysosomal perturbations in human dopaminergic neurons derived from induced pluripotent stem cells with PARK2 mutation. Sci Rep 10, 10278 (2020).

35. Li, K., Li, J., Zheng, J. & Qin, S. Reactive Astrocytes in Neurodegenerative Diseases. Aging Dis 10, 664–675 (2019).

36. Shang, X. et al. Rational design of small molecule inhibitors targeting RhoA subfamily Rho GTPases. Chem Biol 19, 699–710 (2012).

37. Hall, A. & Lalli, G. Rho and Ras GTPases in axon growth, guidance, and branching. Cold Spring Harb Perspect Biol 2, a001818 (2010).

38. Pahan, K. Lipid-lowering drugs. Cell Mol Life Sci 63, 1165–1178 (2006).

39. Cole, S.L. et al. Statins cause intracellular accumulation of amyloid precursor protein, beta-secretase-cleaved fragments, and amyloid beta-peptide via an isoprenoid-dependent mechanism. J Biol Chem 280, 18755–18770 (2005).

40. Cordle, A. & Landreth, G. 3-Hydroxy-3-methylglutaryl-coenzyme A reductase inhibitors attenuate beta-amyloid-induced microglial inflammatory responses. J Neurosci 25, 299–307 (2005).

41. Ng, M.Y.W., Wai, T. & Simonsen, A. Quality control of the mitochondrion. Dev Cell 56, 881–905 (2021).

42. Tian, R. et al. CRISPR Interference-Based Platform for Multimodal Genetic Screens in Human iPSC-Derived Neurons. Neuron 104, 239–255 e212 (2019).

43. Saeedi Saravi, S.S., Saeedi Saravi, S.S., Khoshbin, K. & Dehpour, A.R. Current insights into pathogenesis of Parkinson’s disease: Approach to mevalonate pathway and protective role of statins. Biomed Pharmacother 90, 724–730 (2017).

44. Mady, A. et al. Determining the role of statins in Parkinson’s disease risk reduction and disease modification: A comprehensive meta-analysis of 4 million participants’ data. CNS Neurosci Ther 30, e14888 (2024).

45. Wolozin, B. et al. Simvastatin is associated with a reduced incidence of dementia and Parkinson’s disease. BMC Med 5, 20 (2007).

46. Gao, X., Simon, K.C., Schwarzschild, M.A. & Ascherio, A. Prospective study of statin use and risk of Parkinson disease. Arch Neurol 69, 380–384 (2012).

47. Ritz, B. et al. Statin use and Parkinson’s disease in Denmark. Mov Disord 25, 1210–1216 (2010).

48. Becker, C., Jick, S.S. & Meier, C.R. Use of statins and the risk of Parkinson’s disease: a retrospective case-control study in the UK. Drug Saf 31, 399–407 (2008).

49. Samii, A., Carleton, B.C. & Etminan, M. Statin use and the risk of Parkinson disease: a nested case control study. J Clin Neurosci 15, 1272–1273 (2008).

50. Huang, X. et al. Statins, plasma cholesterol, and risk of Parkinson’s disease: a prospective study. Mov Disord 30, 552–559 (2015).

51. Liu, G. et al. Statins may facilitate Parkinson’s disease: Insight gained from a large, national claims database. Mov Disord 32, 913–917 (2017).

52. Goncalves, O.R. et al. Statins as potential disease-modifying therapy of Parkinson’s disease: a systematic review and meta-analysis with trial sequential analysis. Acta Neurol Belg 125, 915–922 (2025).

53. Stevens, K.N. et al. Evaluation of Simvastatin as a Disease-Modifying Treatment for Patients With Parkinson Disease: A Randomized Clinical Trial. JAMA Neurol (2022).

54. Lin, C.H. et al. A Double-Blind, Randomized, Controlled Trial of Lovastatin in Early-Stage Parkinson’s Disease. Mov Disord 36, 1229–1237 (2021).

55. Tong, H. et al. Simvastatin Inhibits Activation of NADPH Oxidase/p38 MAPK Pathway and Enhances Expression of Antioxidant Protein in Parkinson Disease Models. Front Mol Neurosci 11, 165 (2018).

56. Bar-On, P. et al. Statins reduce neuronal alpha-synuclein aggregation in in vitro models of Parkinson’s disease. J Neurochem 105, 1656–1667 (2008).

57. Xu, Y.Q. et al. Simvastatin induces neuroprotection in 6-OHDA-lesioned PC12 via the PI3K/AKT/caspase 3 pathway and anti-inflammatory responses. CNS Neurosci Ther 19, 170–177 (2013).

58. Ghosh, A. et al. Simvastatin inhibits the activation of p21ras and prevents the loss of dopaminergic neurons in a mouse model of Parkinson’s disease. J Neurosci 29, 13543–13556 (2009).

59. Ostrowski, S.M., Wilkinson, B.L., Golde, T.E. & Landreth, G. Statins reduce amyloid-beta production through inhibition of protein isoprenylation. J Biol Chem 282, 26832–26844 (2007).

60. Kim, W.Y. et al. Statins decrease dendritic arborization in rat sympathetic neurons by blocking RhoA activation. J Neurochem 108, 1057–1071 (2009).

61. Hamano, T., Yen, S.H., Gendron, T., Ko, L.W. & Kuriyama, M. Pitavastatin decreases tau levels via the inactivation of Rho/ROCK. Neurobiol Aging 33, 2306–2320 (2012).

62. Pooler, A.M., Xi, S.C. & Wurtman, R.J. The 3-hydroxy-3-methylglutaryl co-enzyme A reductase inhibitor pravastatin enhances neurite outgrowth in hippocampal neurons. J Neurochem 97, 716–723 (2006).

63. Lopez-Lopez, A., Labandeira, C.M., Labandeira-Garcia, J.L. & Munoz, A. Rho kinase inhibitor fasudil reduces l-DOPA-induced dyskinesia in a rat model of Parkinson’s disease. Br J Pharmacol 177, 5622–5641 (2020).

64. Cartocci, V. et al. Modulation of the Isoprenoid/Cholesterol Biosynthetic Pathway During Neuronal Differentiation In Vitro. J Cell Biochem 117, 2036–2044 (2016).

65. Galiullina, L.F., et al. Interaction of statins with phospholipid bilayers studied by solid-state NMR spectroscopy. Biochim Biophys Acta Biomembr 1861, 584–593 (2019).

66. Khodov, I.A., Huster, D. & Scheidt, H.A. The interaction of small molecules with phospholipid membranes studied by solid-state NMR and molecular dynamics simulation. Biophys Rev 17, 1401–1413 (2025).

67. Sanchez-Danes, A. et al. Disease-specific phenotypes in dopamine neurons from human iPS-based models of genetic and sporadic Parkinson’s disease. EMBO Mol Med 4, 380–395 (2012).

68. Borgs, L. et al. Dopaminergic neurons differentiating from LRRK2 G2019S induced pluripotent stem cells show early neuritic branching defects. Sci Rep 6, 33377 (2016).

69. Gcwensa, N.Z., Russell, D.L., Cowell, R.M. & Volpicelli-Daley, L.A. Molecular Mechanisms Underlying Synaptic and Axon Degeneration in Parkinson’s Disease. Front Cell Neurosci 15, 626128 (2021).

70. Yang, L., Mao, K., Yu, H. & Chen, J. Neuroinflammatory Responses and Parkinson’ Disease: Pathogenic Mechanisms and Therapeutic Targets. J Neuroimmune Pharmacol 15, 830–837 (2020).

71. MacMahon Copas, A.N., McComish, S.F., Fletcher, J.M. & Caldwell, M.A. The Pathogenesis of Parkinson’s Disease: A Complex Interplay Between Astrocytes, Microglia, and T Lymphocytes? Front Neurol 12, 666737 (2021).

72. Menassa, D.A. & Gomez-Nicola, D. Microglial Dynamics During Human Brain Development. Front Immunol 9, 1014 (2018).

73. Koch, J.C. et al. ROCK inhibition in models of neurodegeneration and its potential for clinical translation. Pharmacol Ther 189, 1–21 (2018).

74. Lau, C.L. et al. Transcriptomic profiling of astrocytes treated with the Rho kinase inhibitor fasudil reveals cytoskeletal and pro-survival responses. J Cell Physiol 227, 1199–1211 (2012).

75. Du, R.W. & Bu, W.G. Simvastatin Prevents Neurodegeneration in the MPTP Mouse Model of Parkinson’s Disease via Inhibition of A1 Reactive Astrocytes. Neuroimmunomodulation 28, 82–89 (2021).

76. Kumar, A., Sharma, N., Gupta, A., Kalonia, H. & Mishra, J. Neuroprotective potential of atorvastatin and simvastatin (HMG-CoA reductase inhibitors) against 6-hydroxydopamine (6-OHDA) induced Parkinson-like symptoms. Brain Res 1471, 13–22 (2012).

77. Hirsch, E.C. & Hunot, S. Neuroinflammation in Parkinson’s disease: a target for neuroprotection? Lancet Neurol 8, 382–397 (2009).

78. Ransohoff, R.M. How neuroinflammation contributes to neurodegeneration. Science 353, 777–783 (2016).

79. Ryan, B.J., Hoek, S., Fon, E.A. & Wade-Martins, R. Mitochondrial dysfunction and mitophagy in Parkinson’s: from familial to sporadic disease. Trends Biochem Sci 40, 200–210 (2015).

80. Ahmed, L.A., Darwish, H.A., Abdelsalam, R.M. & Amin, H.A. Role of Rho Kinase Inhibition in the Protective Effect of Fasudil and Simvastatin Against 3-Nitropropionic Acid-Induced Striatal Neurodegeneration and Mitochondrial Dysfunction in Rats. Mol Neurobiol 53, 3927–3938 (2016).

81. Paintlia, A.S., Paintlia, M.K., Singh, A.K. & Singh, I. Modulation of Rho-Rock signaling pathway protects oligodendrocytes against cytokine toxicity via PPAR-alpha-dependent mechanism. Glia 61, 1500–1517 (2013).

82. Carroll, C.B. & Wyse, R.K.H. Simvastatin as a Potential Disease-Modifying Therapy for Patients with Parkinson’s Disease: Rationale for Clinical Trial, and Current Progress. J Parkinsons Dis 7, 545–568 (2017).

83. Gegg, M.E. & Schapira, A.H. Mitochondrial dysfunction associated with glucocerebrosidase deficiency. Neurobiol Dis 90, 43–50 (2016).

84. Fernandez-Mosquera, L. et al. Acute and chronic mitochondrial respiratory chain deficiency differentially regulate lysosomal biogenesis. Sci Rep 7, 45076 (2017).

85. Ivankovic, D., Chau, K.Y., Schapira, A.H. & Gegg, M.E. Mitochondrial and lysosomal biogenesis are activated following PINK1/parkin-mediated mitophagy. J Neurochem 136, 388–402 (2016).

86. Zhang, X., Zhang, H., Dong, J., Cai, H. & Le, W. Advances in autophagy for Parkinson’s disease pathogenesis and treatment. Ageing Neurodegener Dis 5 (2025).

87. Zhang, Y. et al. Simvastatin improves lysosome function via enhancing lysosome biogenesis in endothelial cells. Front Biosci (Landmark Ed) 25, 283–298 (2020).

88. Moskal, N. et al. ROCK inhibitors upregulate the neuroprotective Parkin-mediated mitophagy pathway. Nat Commun 11, 88 (2020).

89. Clark, E.H., Vazquez de la Torre, A., Hoshikawa, T. & Briston, T. Targeting mitophagy in Parkinson’s disease. J Biol Chem 296, 100209 (2021).

90. Kang, S.Y. et al. Autophagic modulation by rosuvastatin prevents rotenone-induced neurotoxicity in an in vitro model of Parkinson’s disease. Neurosci Lett 642, 20–26 (2017).

91. Wang, H. et al. Tumorous cholesterol biosynthesis curtails anti-tumor immunity by preventing MTOR-TFEB-mediated lysosomal degradation of CD274/PD-L1. Autophagy 21, 2670–2689 (2025).

92. Xu, X. et al. Simvastatin promotes NPC1-mediated free cholesterol efflux from lysosomes through CYP7A1/LXRα signalling pathway in oxLDL-loaded macrophages. J Cell Mol Med 21, 364–374 (2017).

93. Davis, O.B. et al. NPC1-mTORC1 Signaling Couples Cholesterol Sensing to Organelle Homeostasis and Is a Targetable Pathway in Niemann-Pick Type C. Dev Cell 56, 260–276.e7 (2021).

94. Mattson, M.P., Gleichmann, M. & Cheng, A. Mitochondria in neuroplasticity and neurological disorders. Neuron 60, 748–766 (2008).

95. Ni, A. & Ernst, C. Evidence That Substantia Nigra Pars Compacta Dopaminergic Neurons Are Selectively Vulnerable to Oxidative Stress Because They Are Highly Metabolically Active. Front Cell Neurosci 16, 826193 (2022).

96. Kast, D.J. & Dominguez, R. The Cytoskeleton-Autophagy Connection. Curr Biol 27, R318–R326 (2017).

97. Nishimura, Y., Itoh, K., Yoshioka, K., Ikeda, K. & Himeno, M. A role for small GTPase RhoA in regulating intracellular membrane traffic of lysosomes in invasive rat hepatoma cells. Histochem J 34, 189–213 (2002).

