## Supplementary Information for "Simvastatin attenuates disease phenotypes in human induced pluripotent stem cell models of familial Parkinson’s disease through RhoA inhibition"

#### Product code list

| Reagent | Company | Catalogue number |
| --- | --- | --- |
| **Isogenic iPSC derived NSC maintenance and differentiation** |  |  |
| Geltrex | Thermo Fischer | A1413202 |
| Neurobasal medium | Thermo Fischer | 21103049 |
| Non-essential amino acids | Thermo Fischer | 11140050 |
| GlutaMAX-1 | Thermo Fischer | 35050038 |
| B-27 Supplement | Thermo Fischer | 17504044 |
| Penicillin-streptomycin | Sigma-Aldrich | 15140122 |
| bFGF | Sigma-Aldrich | F3685 |
| Accutase | Thermo Fischer | A1110501 |
| Poly-L-ornithine | Sigma-Aldrich | P3655 |
| Laminin | Thermo Fischer | 23017015 |
| Dopaminergic Induction Pack | XCell Science | DI-001 |
| Dopaminergic Maturation Pack | XCell Science | DM-001 |
| Sonic Hedgehog | Peprtech | 100-45 |
| **PD patient iPSC maintenance and differentiation** |  |  |
| Geltrex | Thermo Fischer | A1413202 |
| mTeSR1 | Stem Cell Technologies | 85850 |
| TrypLE | Thermo Fischer | 126040 |
| ROCK inhibitor Y27632 | Bio-Techne | 1254 |
| Knockout DMEM | Thermo Fischer | 10829018 |
| KnockOut serum replacement | Thermo Fischer | 10828010 |
| β-mercaptoethanol | Sigma-Aldrich | M6250 |
| L-glutamine | Thermo Fischer | 25030-032 |
| N2 supplement | Thermo Fischer | 17502048 |
| B-27 Supplement, minus vitamin A | Thermo Fischer | 12587010 |
| LDN193189 | Sigma-Aldrich | SML0559 |
| SB431542 | Bio-Techne | 1614 |
| Sonic Hedgehog-C24II | Bio-Techne | 1845-SH |
| Purmorphamine | Bio-Techne | 4551 |
| FGF8 | Strattech | 16124-HNAE-SIB |
| CHIR99021 | Bio-Techne | 4423 |
| Brain-derived neurotrophic factor | Peprotech | 450-02 |
| Glial cell line-derived neurotrophic factor | Peprotech | 450-10 |
| Ascorbic acid | Sigma-Aldrich | A4544 |
| Transforming growth factor type $\beta 3$ | Peprotech | 10036E |
| Dibutyl cAMP | Sigma-Aldrich | D0627 |
| DAPT | Abcam | 120633 |
| Poly-L-ornithine | Sigma | P4957 |
| Biolaminin | BioLamina | LN521 |
| Antibiotic Antimycotic | Sigma-Aldrich | A5955 |
| Mitomycin C | Sigma-Aldrich | M4287 |
| Neurobasal medium, minus phenol red | Thermo Fischer | 12348017 |
| **Treatments** |  |  |
| Rhosin | Tocris | 5003 |
| Simvastatin | Sigma-Aldrich | S6196 |
| Gemfibrozil | Sigma-Aldrich | G9518 |
| Rho inhibitor I (C3 transferase) | Cytoskeleton Inc. | CT04 |
| Carbonyl cyanide 3-chlorophenylhydrazone (CCCP) | Sigma-Aldrich | C2759 |
| Depheriprone | Cayman Chemical | 20387 |
| **Immunocytochemistry** |  |  |
| Paraformaldehyde | Sigma-Aldrich | 158127 |
| D-PBS | Thermo Fischer | 14190144 |
| Triton X-100 | Sigma-Aldrich | 11332481001 |
| Saponin | Sigma-Aldrich | 84510 |
| Donkey serum | Merck Millipore | S30 |
| Goat serum | Merck Millipore | S26 |
| Fetal bovine serum | Thermo Fischer | 10082147 |
| DAPI | Sigma-Aldrich | D9542 |
| ProLong® Diamond mounting medium | Thermo Fischer | P36961 |
| Filipin | Sigma-Aldrich | F9765 |
| HRP-conjugated streptavidin | Sigma-Aldrich | GERPN1231 |
| 3,3'-diaminobenzidine | Sigma-Aldrich | D5637 |
| H<sub>2</sub>O<sub>2</sub> | Merck Millipore | 107209 |
| **Western blotting** |  |  |
| RIPA | Thermo Fischer | 89900 |
| cOmplete™, Mini, EDTA-free Protease Inhibitor Cocktail | Sigma-Aldrich | 11836170001 |
| Phosphatase inhibitor (PhosSTOP tablets) | Sigma-Aldrich | 4906845001 |
| Bicinchoninic acid assay (BCA) | Pierce | 23225 |
| NuPAGE LDS Sample Buffer | Thermo Fischer | NP0007 |
| 4-12% Bis-Tris gels | Thermo Fischer | NP0335 |
| NuPAGE MOPS SDS Running Buffer | Thermo Fischer | NP0001 |
| SeeBlue™ Plus2 Pre-stained Protein Standard | Thermo Fischer | LC5925 |
| NuPAGE Antioxidant | Thermo Fischer | NP0005 |
| iBlot™ Transfer Stack, PVDF, mini | Thermo Fischer | IB401002 |
| Trizma base | Sigma-Aldrich | T1378 |
| NaCl | Sigma-Aldrich | 31434 |
| HCl | VWR | 20252 |
| Tween-20 | Sigma-Aldrich | P1379 |
| Skim milk | Natur Drogeriet | 32711009 |
| Bovine serum albumin | Sigma-Aldrich | A4503 |
| Pierce™ ECL Western Blotting Substrate | Thermo Fischer | 32109 |
| **Antibodies** |  |  |
| Rabbit anti-tyrosine hydroxylase | Merck Millipore | AB152 |
| Sheep anti-tyrosine hydroxylase | Merck Millipore | AB1542 |
| Goat anti-FOXA2 | Bio-Techne | AF2400 |
| Rabbit anti-LMX1A | Abcam | ab139726 |
| Rabbit anti-β-tubulin III | Sigma-Aldrich | T2200 |
| Mouse anti-MAP2a+b | Sigma-Aldrich | M1406 |
| Chicken anti-MAP2a+b+c | Merck Millipore | AB5543 |
| Mouse anti-synaptophysin | Merck Millipore | S5768 |
| Guiney pig anti-synapsin | Synaptic systems | 106004 |
| Rabbit anti-homer 1 | Synaptic systems | 160003 |
| Rabbit anti-glial fibrillary acidic protein | DAKO | Z0334 |
| Rabbit anti-TOM20 | Santa-Cruz | SC11415 |
| Mouse anti-TOM20 | Santa-Cruz | SC17764 |
| Rabbit anti-LAMP1 | Abcam | 108597 |
| Mouse anti-LAMP1 | Santa-Cruz | SC20011 |
| Rabbit anti-cleaved caspase 3 | Cell Signalling | 9661 |
| Rabbit anti-NIX | Abcam | AB8399 |
| Rabbit anti-NIX | Cell Signalling | 12396 |
| Mouse anti-Parkin | Cell Signalling | 4211 |
| Mouse anti-P62/SQSTM1 | Abcam | 56416 |
| Rabbit anti-LC3B | Cell Signalling | 3868 |
| Mouse anti-RhoA | Santa-Cruz | SC418 |
| Rabbit anti-ATP<sub>ase</sub> α-2 | Merck Millipore | 07-674 |
| Mouse anti-actin | Merck Millipore | MAB1501 |
| Mouse anti-human nuclei | Merck Millipore | MAB1281 |
| Goat anti-rabbit AF488 | Thermo Fischer | A11008 |
| Goat anti-mouse AF488 | Thermo Fischer | A11001 |
| Goat anti-guinea pig AF488 | Abcam | Ab96459 |
| Donkey anti-rabbit AF488 | Thermo Fischer | A21206 |
| Donkey anti-mouse AF488 | Thermo Fischer | A21202 |
| Donkey anti-goat AF488 | Thermo Fischer | A11055 |
| Donkey anti-sheep AF488 | Thermo Fischer | A11015 |
| Goat anti-rabbit AF555 | Thermo Fischer | A21428 |
| Goat anti-mouse AF555 | Thermo Fischer | A21422 |
| Donkey anti-rabbit AF555 | Thermo Fischer | A31572 |
| Donkey anti-mouse AF555 | Thermo Fischer | A31570 |
| Goat anti-chicken AF647 | Thermo Fischer | A21449 |
| Donkey anti-rabbit AF647 | Thermo Fischer | A31573 |
| Donkey anti-sheep AF647 | Thermo Fischer | A21448 |
| HRP-conjugated goat anti-mouse | Agilent | P0447 |
| HRP-conjugated goat anti-rabbit | Agilent | P0448 |
| Biotinylated donkey anti-rabbit IgG | Cytiva | RPN1004 |
| Biotinylated sheep anti-mouse IgG | Cytiva | RPN1001 |
| **Assays** |  |  |
| G-LISA RhoA Activation Assay Biochem Kit | Cytoskeleton Inc. | BK124 |
| Total RhoA ELISA kit | Cytoskeleton Inc. | BK150 |
| CytoTox96® Non-Radioactive Cytotoxicity Assay | Promega | G1780 |
| V-PLEX Proinflammatory Panel 1 Human kit | MesoScale Discovery | K15049D-2 |
| R-PLEX Human Neurofilament L Assay | MesoScale Discovery | K1517XR-2 |
| Fluorometric β-Galactosidase Activity Assay Kit | BioVision | K821 |
| JC-10 | AAT Bioquest®, Inc. | 22204 |
| CellROX® Green Reagent | Molecular Probes | C10444 |
| MitoSOX™ Mitochondrial Superoxide Indicator | Thermo Fischer | M36008 |
| ATPlite Assay | Perkin Elmer | 6016943 |
| **Calcium imaging** |  |  |
| Fluo-4-AM | Thermo Fischer | F14201 |
| Perfusion chamber | Warner instruments | RC-25F |
| KCl | Merck | TA630136 |
| NaCl | Sigma-Aldrich | 31434 |
| MgCl<sub>2</sub> | Merck | 105833 |
| CaCl<sub>2</sub> | Merck | 102382 |
| Hepes | Sigma-Aldrich | H3375 |
| Glucose | Merck | 108342 |

**Supplementary table 1.** Cell line information for the cell lines included in this study.

| Isogenic cell lines |  |  |  |  |  |
| --- | --- | --- | --- | --- | --- |
| Cell line | Cell line ID | Cell type | Sex | Genotype |  |
| Control | XCL-1 | iPSC-derived NSC | Male | wt |  |
| PARK2-/- | PARK2 | iPSC-derived NSC | Male | delEx2; homo |  |
| Parkinson's disease patient-derived lines |  |  |  |  |  |
| Cell line | Cell line ID | Cell type | Sex | Age | Genotype |
| Control 1 | SFC067-03-01 | iPSC | Male | 72 | wt |
| Control 2 | SFC156-03-01 | iPSC | Male | 75 | wt |
| Control 3 | SFC856-03-04 | iPSC | Female | 78 | wt |
| Control 4 | SFC065-03-03 | iPSC | Male | 65 | wt |
| Control 5 | JR053-1 | iPSC | Male | 68 | wt |
| PARKIN 1 | SFC821-03-01 | iPSC | Female | 38 | c.823C>T; c.1054T>C |
| PARKIN 2 | SFC818-03-01 | iPSC | Male | 63 | delEx4 +c.924C>T |
| PARKIN 3 | SFC817-03-04 | iPSC | Male | 82 | c.1072Tdel +delEx7 |
| PINK 1 | SFC826-04-06 | iPSC | Female | 47 | c.1366C<T; homo |
| PINK 2 | SFC825-04-02 | iPSC | Female | 53 | c.1366C<T; homo |
| PINK 3 | SFC824-03-02 | iPSC | Male | 38 | c.1366C<T; homo |
| PINK 4 | SFC823-03-07 | iPSC | Female | 76 | c.1366C>T; homo |
| PINK 5 | SFC822-03-05 | iPSC | Female | 75 | c.509T>G; homo |
| LRRK2 (G2019S) 1 | SFC832-03-06 | iPSC | Female | 77 | G2019S |
| LRRK2 (G2019S) 2 | MK002-4 | iPSC | Female | 72 | G2019S |
| LRRK2 (G2019S) 3 | MK114-7 | iPSC | Female | 57 | G2019S |
| LRRK2 (G2019S) 4 | JR36-1 | iPSC | Male | 49 | G2019S |
| LRKK2 (R1441C) 1 | SFC870-03-05 | iPSC | Male | 32 | R1441C |
| LRKK2 (R1441C) 2 | SFC869-03-01 | iPSC | Female | 56 | R1441C |
| LRKK2 (R1441C) 3 | JR207-1 | iPSC | Female | 72 | R1441C |
| GBA (L444P) 1 | SFC866-03-02 | iPSC | Male | 61 | L444P |
| GBA (L444P) 2 | SFC872-03-02 | iPSC | Male | 68 | L444P |
| GBA (L444P) 3 | JR065-1 | iPSC | Female | 46 | L444P |
| GBA (L444P) 4 | NH031-1 | iPSC | Female | 50 | L444P |
| GBA (N370S) 1 | MK088-1 | iPSC | Male | 46 | N370S |
| GBA (N370S) 2 | MK082-26 | iPSC | Male | 51 | N370S |
| SNCA (A53T) 1 | SFC829-03-02 | iPSC | Male | 46 | A53T |
| SNCA (A53T) 2 | SFC828-03-04 | iPSC | Female | 51 | A53T |
| SNCA (trip.) 1 | SFC831-03-03 | iPSC | Female | 55 | Triplication |
| SNCA (trip.) 2 | SFC831-03-01 | iPSC | Female | 55 | Triplication |
| SNCA (trip.) 3 | SFC831-03-05 | iPSC | Female | 55 | Triplication |
| CRISPRi cell line |  |  |  |  |  |
| Cell line | Cell line ID | Cell type | Sex | Age | Genotype |
| WTC11 | WTC11 | iPSC | Male | 34 | dCas9 |

### Supplementary methods

#### Calcium imaging

Cells were grown on 14 mm coverslips and differentiated until day 25. Cells were washed twice in Krebs-Ringer solution (139 mM NaCl, 5 mM KCl, 1.2 mM MgCl<sub>2</sub>, 2 mM CaCl<sub>2</sub>, 10 mM Hepes, and 10 mM glucose) and incubated for 30 min with 5 μM Fluo-4-AM (Thermo Fischer) at 37°C in the dark. Following fluorophore loading, cells were washed twice in Krebs-Ringer solution, and the coverslip was placed in a perfusion chamber (Warner instruments) in Krebs-Ringer solution. Live cell imaging was performed using the FluoView FV1000MPE multiphoton laser confocal microscope (Olympus), and Fluo-4-AM was excited at 488 nm; emission was measured at 510 nm through a long pass filter. Movies were recorded from 300 frames with 1.6 sec frame interval (∼ 0.6 Hz), and cells were depolarized with 25 mM KCl. Depolarization was performed after the first 100 frames to record the baseline.

Movie analysis was performed using custom-made scripts in MatLab (version R2023a, The Mathworks, Natick, USA) based on the image processing toolbox. The image 150 of the movie sequence was used to detect the individual cells across the entire movie, since this image presented the high stimulation across all cells with no artifacts from the stimulation occurring at image 100. Cell somas from image 150 were detected by isolating circular eroded areas using grayscale. A subsequent filter extracted all connected pixels in that area from circular shapes across the images. The filtered image was then dilated to extract the correct cells, which were finally labelled as individual cells based on the connectivity of the dilated components of the image. Fluorescence intensity over time was examined for each cell soma in the movies (150-200 cells/movie) and reported as the mean value per frame. Background fluorescence was subtracted by applying the rolling ball background subtraction method, followed by a curve smoothing using a 5-frame moving average. Fluorescence measurements for the individual cells were expressed as the ratio ($\Delta F/F_0$) of mean change in fluorescence ($\Delta F = F - F_0$) relative to the baseline fluorescence ($F_0$). Baseline subtraction is necessary to account for differences in fluorophore loading between cells, as well as the intrinsic increase in fluorescence in calcium imaging. Finally, the $\Delta F/F_0$ curves were normalized to the mean peak value of control cultures for each experiment, to account for signal differences between independent experiments. The mean of normalized $\Delta F/F_0$ values were plotted over time (presented as frame numbers) from 3 movies per independent experiment.

#### **DAB staining**

Fixation and incubation with primary antibodies were performed as described for immunofluorescence staining. The following primary antibodies and dilutions were applied: rabbit anti-tyrosine hydroxylase (TH, Millipore) 1:600 and mouse anti-human nuclei (HN, Millipore) 1:500. Following incubation with primary antibodies ON at 4°C, cells were washed three times in 0.05 M TBS/0.1% Triton X for 15 min at RT and incubated with biotinylated secondary antibodies, diluted in blocking buffer, for 2 hours at RT. The following secondary antibodies and dilutions were applied: biotinylated donkey anti-rabbit IgG (Cytiva) 1:200 and biotinylated sheep anti-mouse IgG (Cytiva) 1:200.

Cells were washed three times in 0.05 M TBS/0.1% Triton X for 15 min at RT and incubated 1 hour at RT with horseradish peroxidase (HRP) conjugated streptavidin (Sigma) diluted 1:200 in TBS/10% fetal bovine serum (FBS, ThermoFisher). Cells were washed three times for 15 min in 0.05 M TBS before visualization with 0.01% 3,3'-diaminobenzidine (DAB, Sigma) and 0.015% H<sub>2</sub>O<sub>2</sub> (Merck) in TBS for 20 min. Cells were washed three times for 15 min in 0.05 M TBS, and imaging was performed using an inverted bright-field microscope (Olympus).

#### **Filipin staining**

Intracellular cholesterol levels were quantified using filipin staining. Neuronal cultures were fixed in 4% paraformaldehyde (PFA, Sigma) in 0.1 M PBS for 15 min at RT and washed twice in PBS. PFA was quenched with 50 mM glycine diluted in PBS before 2 hours incubation at RT with 50 µg/ml filipin (Sigma) diluted in M1 medium containing 150 mM NaCl, 5 mM KCl, 1mM CaCl<sub>2</sub>, 1 mM MgCl<sub>2</sub>, 5 mM glucose, and 20 mM HEPES (pH 7.4). Cells were subsequently washed twice in M1 medium, and coverslips were mounted on glass slides. Filipin staining was imaged with a Leica filter A-cube harbouring a 360 nm bandpass excitation filter, a 400 nm dichromatic filter, and a 425 nm long-pass emission filter. Five randomly chosen fields/coverslip were imaged from independent differentiations. Staining intensity was quantified in ImageJ.

#### **CRISPRi knockdown**

CRISPRi knockdown of *PARK2* was performed in the healthy iPSC cell line WTC11, which constitutively expresses the catalytically dead Cas9 (dCas9) machinery<sup>33</sup>.

WTC11 iPSCs were differentiated using the dual SMAD inhibition protocol as described above. At day 20 of differentiation, the cells were transduced with a lentiviral construct expressing sgRNA targeting *PARK2* or non-targeting (NT).

### Supplementary figures

A

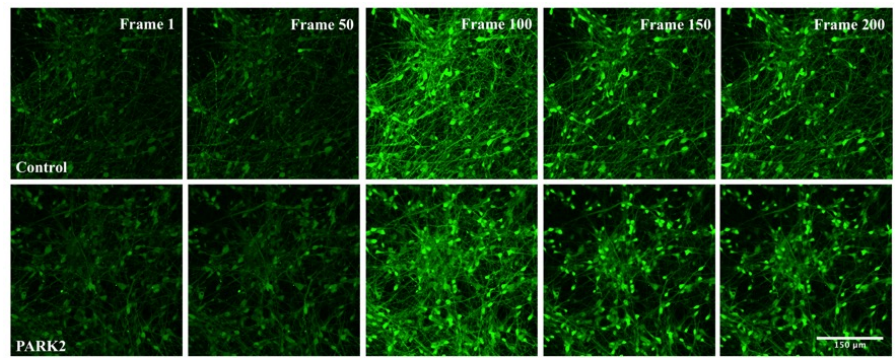

B

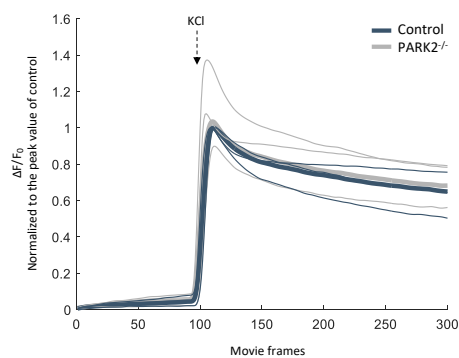

C

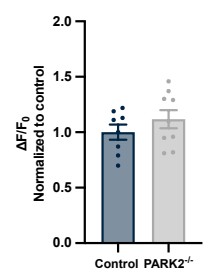

**Suppl. figure 1. Calcium imaging recordings of differentiated *PARK2*<sup>-/-</sup> and isogenic control cultures showing depolarization by potassium chloride (KCl).** A) Movies of Fluo-4-AM loaded *PARK2*<sup>-/-</sup> and isogenic control cultures were recorded from 300 frames with 1.6 sec interval. After the first 100 frames, cells were depolarized with KCl (arrow). The timelapse pictures are shown with 50 frames interval. Scale bar: 150 μm. B) $\Delta F/F_0$ over time (represented by the number of movie frames) was calculated and normalized to the peak value of control cultures for each experiment. Three independent experiments were performed, with 3 movies recorded in each experiment containing between 150-200 cells. The mean from each experiment is shown in thin lines. The mean across all experiments is shown in bold. C) A plot of peak $\Delta F/F_0$ values, showing no difference in the depolarization capacity of control vs. *PARK2*<sup>-/-</sup> neuronal cultures. Mean ± SEM, n = 9 movies, from 3 independent differentiations. Statistical analysis: Student's t-test.

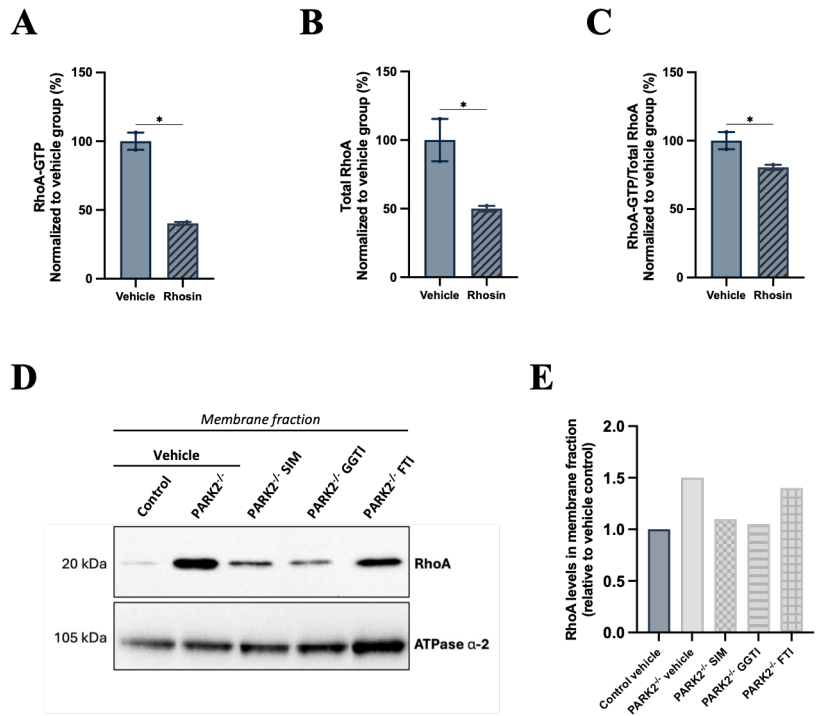

**Suppl. figure 2. RhoA inhibition verification.** A-C) Treatment with 120 μM rhosin for 24 hours reduced RhoA activity as evaluated by the amount of GTP-bound RhoA (control cells). Because the total level of RhoA was reduced with rhosin treatment, the amount of GTP-bound RhoA/total RhoA with rhosin treatment was less pronounced. Mean ± SEM, n = 2 samples per group from 1 differentiation. Statistical analysis: Student's t-test, \*p < 0.05. D-E) Subcellular fractionation revealed increased membrane-associated RhoA in *PARK2*<sup>-/-</sup> cells compared to isogenic control neurons, consistent with increased RhoA-GTP levels in *PARK2*<sup>-/-</sup> cells. Treatment with 300 nM simvastatin (SIM) for 24 hours reduced RhoA membrane association to control levels. Treatment with 5 μM geranylgeranyltransferase inhibitor (GGTI) but not 5 μM farnesyltransferase inhibitor (FTI) for 24 hours mimicked the effect of statins, suggesting that the effect of statins is mediated by inhibition of geranylgeranylation of RhoA required for its membrane association. N = 1 sample per group from 1 differentiation.

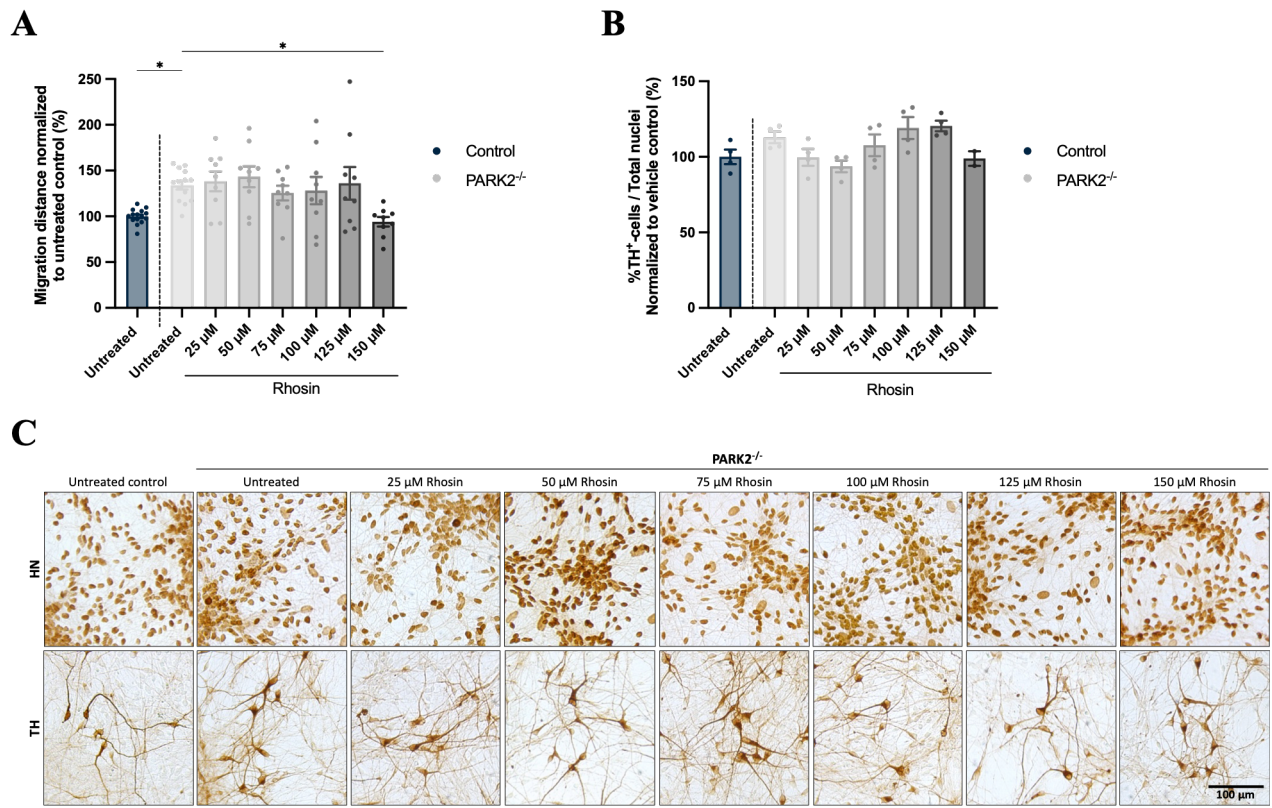

**Suppl. figure 3. Dose-response and toxicity of rhosin treatment on *PARK2*<sup>-/-</sup> cultures.** A) Increasing concentrations of rhosin (25, 50, 75, 100, 125, and 150 μM) were tested for their effect on the abnormal migration of *PARK2*<sup>-/-</sup> neuroblasts. 150 μM rhosin was found most effective in normalizing the migration of *PARK2*<sup>-/-</sup> neuroblasts to a level equal to that of the control. Mean ± SEM, n = 9-15 wells per group from 2-3 independent differentiations. B and C) DAB staining for human nuclei (HN) and the dopaminergic neuronal marker tyrosine hydroxylase (TH) revealed no toxic effects of the concentrations tested. Scale bar: 100 μm. Mean ± SEM, n = 2-4 wells per group from 1 differentiation. Statistical analysis: one-way ANOVA, \*p < 0.05.

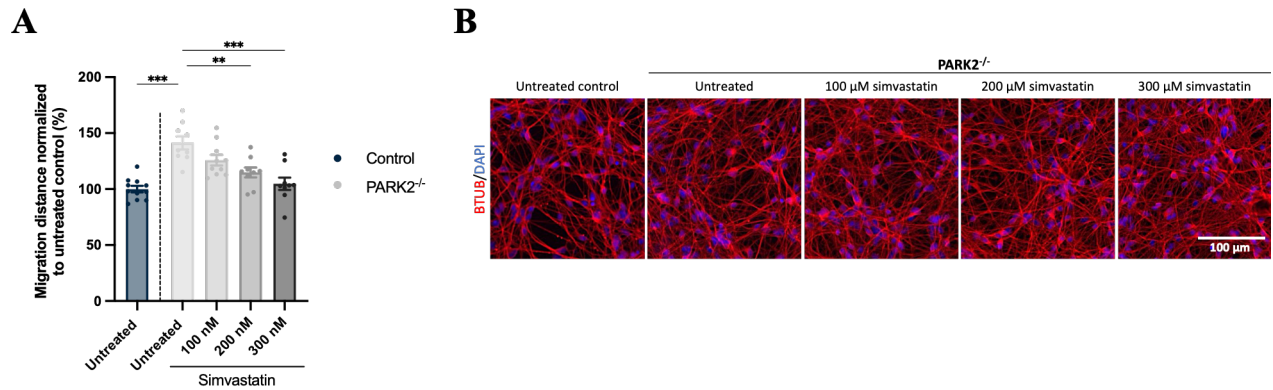

**Suppl. figure 4. Dose-response and toxicity of simvastatin treatment on *PARK2*<sup>-/-</sup> cultures.** A) Increasing concentrations of simvastatin (100, 200, and 300 nM) were tested for their effect on the abnormal migration of *PARK2*<sup>-/-</sup> neuroblasts. 300 nM rhosin was found most effective in normalizing the migration of *PARK2*<sup>-/-</sup> neuroblasts to a level equal to that of the control. Mean ± SEM, n = 9-11 wells per group from 2 independent differentiations. Statistical analysis: one-way ANOVA, \*\*p < 0.01, \*\*\*p < 0.001. B) Immunofluorescence staining for the neuronal marker β-tubulin-III<sup>+</sup> (BTUB) revealed no toxic effects of the concentrations tested. Scale bar: 100 μm.

**A**

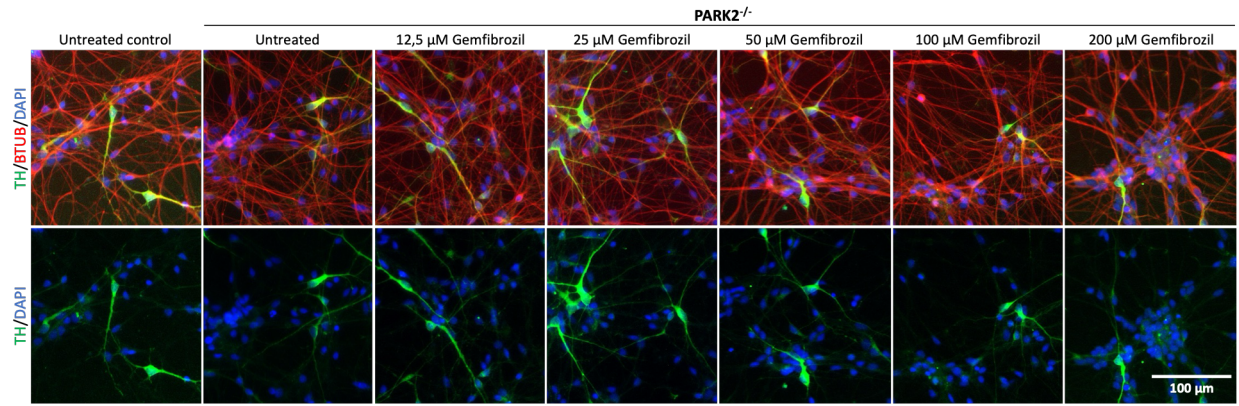

**B**

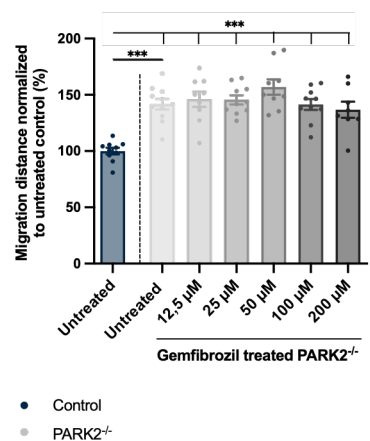

**C**

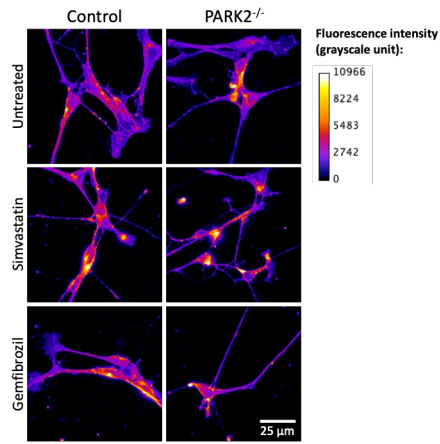

**D**

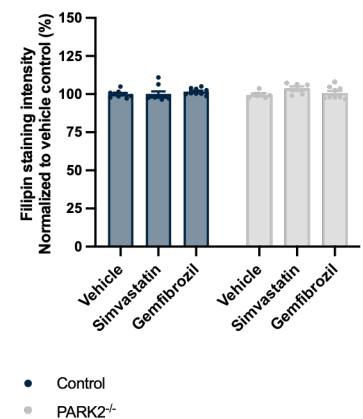

**Suppl. figure 5. Toxicity test of gemfibrozil treatment on *PARK2*<sup>-/-</sup> cultures and treatment effect on cholesterol levels.** A) Increasing concentrations of gemfibrozil (12.5, 25, 50, 100, and 200 µM) were tested for their toxic effect on *PARK2*<sup>-/-</sup> neurons. Immunofluorescence staining for the neuronal marker β-tubulin-III<sup>+</sup> (BTUB, red) and the dopaminergic neuronal marker tyrosine hydroxylase (TH, green) revealed no toxic effects of the concentrations tested. Scale bar: 100 µm. B) Treatment with gemfibrozil did not rescue the increased migration of *PARK2*<sup>-/-</sup> neurons for any of the concentrations tested (12.5 – 200 µM). Mean ± SEM, n = 8-11 wells from 2 independent differentiations. Statistical analysis: one-way ANOVA, \*\*\*p < 0.001. C-D) Quantification of intracellular cholesterol levels by filipin staining showed no reduction in cholesterol by simvastatin (300 nM) and gemfibrozil treatment (50 µM) under the experimental conditions for either *PARK2*<sup>-/-</sup> or isogenic control neurons. Scale bar = 25 µm. Mean ± SEM, n = 6-9 coverslips per group from 2 independent differentiations. Statistical analysis: one-way ANOVA.

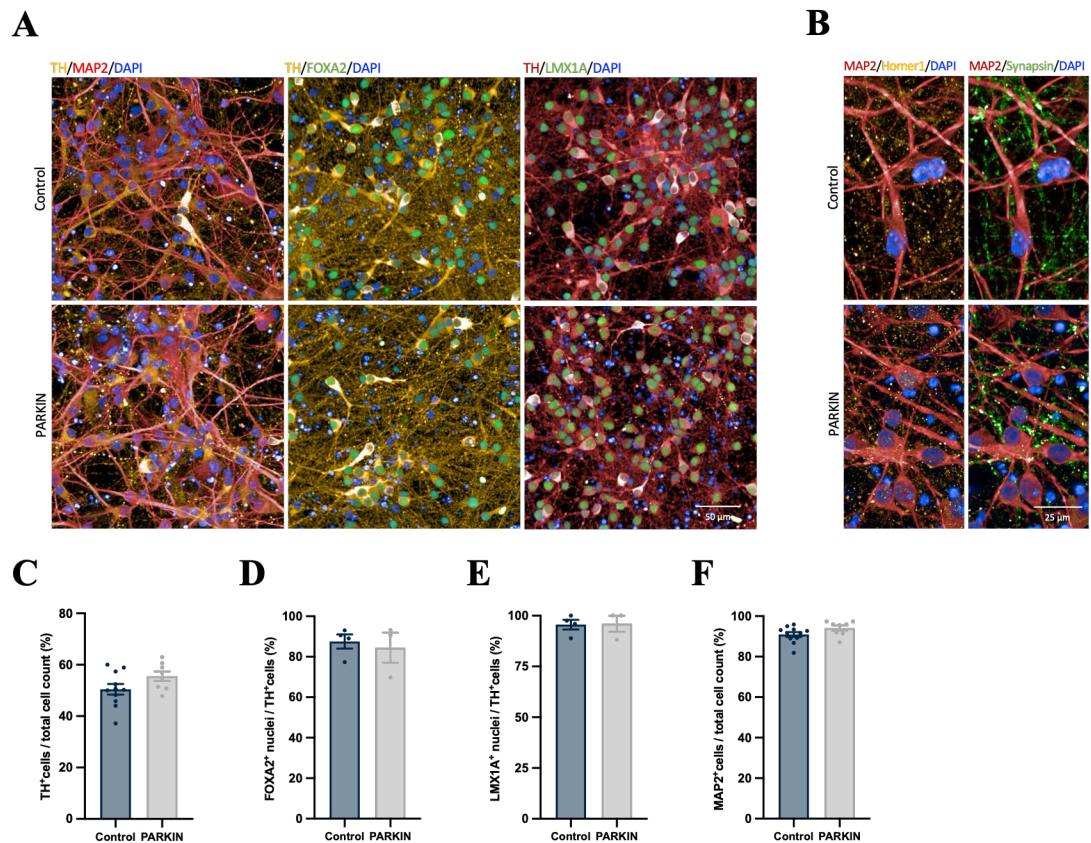

**Suppl. figure 6. General characterization of parkin PD patient iPSC-derived neuronal cultures.**

A-F) PD patient-derived iPSC lines from three *PARK2* mutation carriers and five healthy controls were differentiated into DA neuronal cultures for 35 days. Immunofluorescence staining and quantification of tyrosine hydroxylase (TH, yellow), microtubule-associated protein 2 (MAP2, red), FOXA2 (green), LMX1A (green), homer 1 (yellow), and synapsin 1 (green) showed no difference in the differentiation capacity of parkin and PINK1 PD patient-derived cell lines relative to healthy control cell lines. Mean ± SEM, $n = 8-11$ cell line mean values from 3 independent differentiations (C and F), and $n = 3-4$ cell line mean values from 1 differentiation (D and E). Statistical analysis: student's t-test.

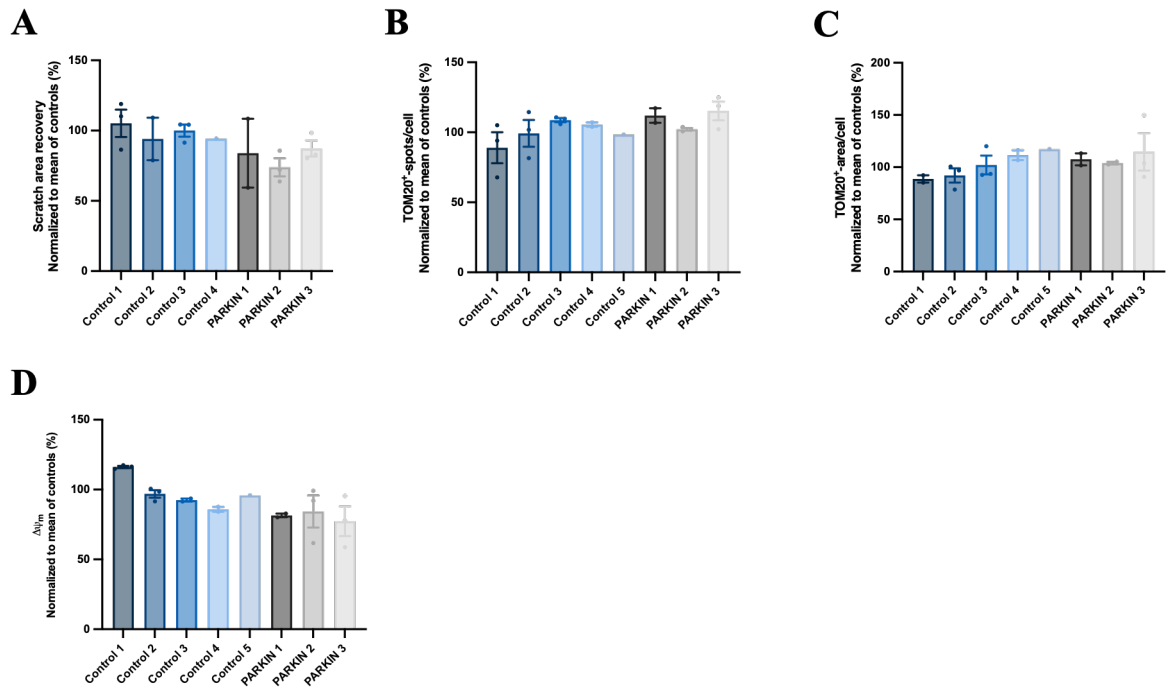

**Suppl. figure 7. Individual cell line data for parkin and healthy control PD patient-derived cell lines.** A) Scratch assay data evaluating neurite outgrowth for the healthy control cell lines and parkin PD patient-derived cell lines included in the study. B-C) Individual cell line data for mitochondrial number and area evaluated from immunofluorescence staining of the outer mitochondrial membrane protein TOM20. D) Mitochondrial membrane potential ($\Delta\psi_m$) for the included cell lines. Mean ± SEM, n = 1-3 differentiations per cell line. No statistical analysis performed.

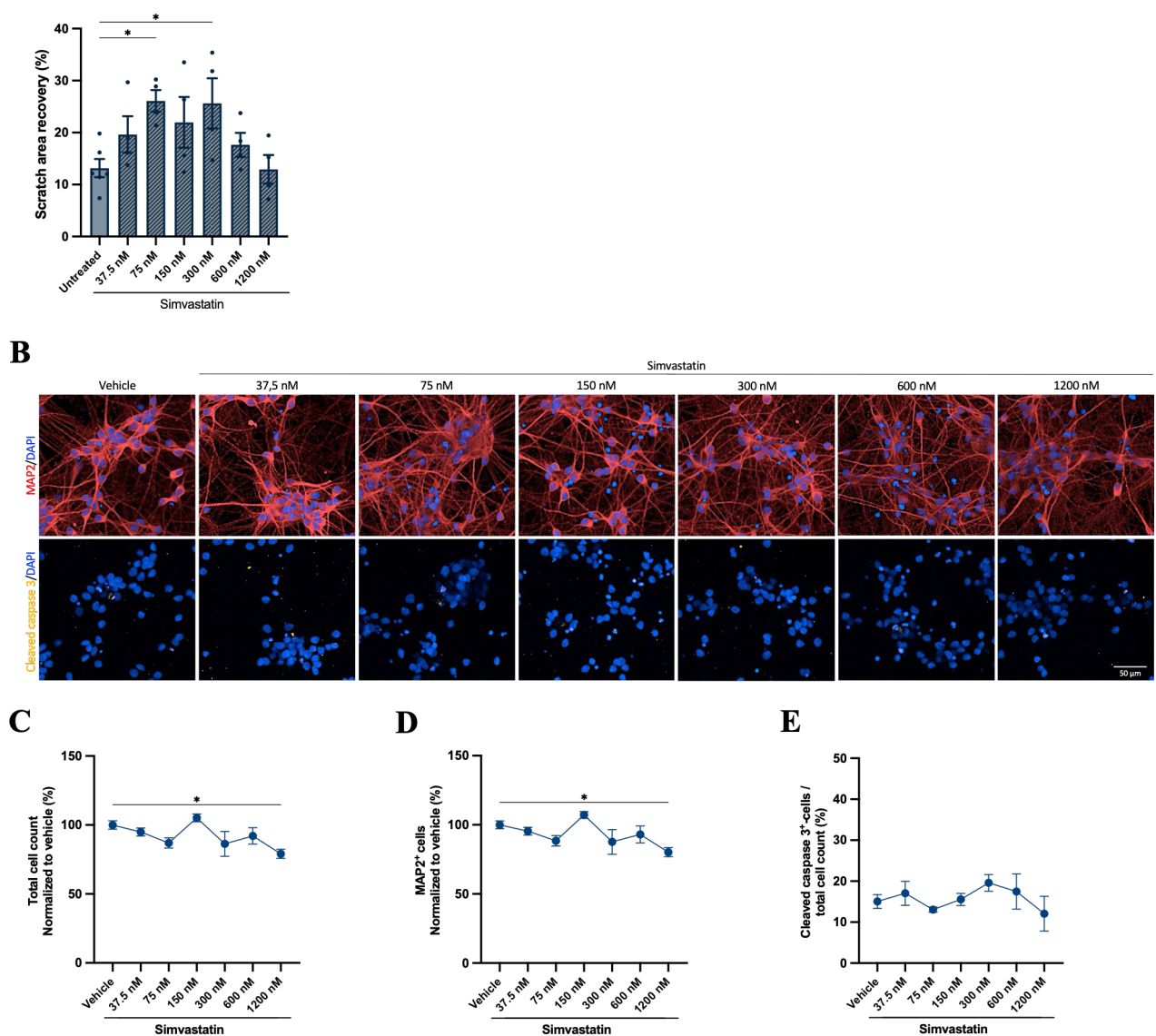

**Suppl. figure 8. Dose-response and toxicity of simvastatin treatment on patient iPSC-derived neurons.** A) Increasing concentrations of simvastatin were tested for their effect on neurite outgrowth of a healthy control patient line. A scratch assay was applied to evaluate the outgrowth of neurites into the scratch area by measuring the area covered by neurites (% scratch area recovery). Mean ± SEM, n = 4-6 wells per group from 1 differentiation. B) Immunofluorescence staining for microtubule-associated protein 2 (MAP2, red) and cleaved caspase 3 (yellow). Scale bar: 50 μm. C-E) Staining quantification revealed no toxic effect of the tested simvastatin concentrations as evaluated by the total cell number calculated from the number of DAPI<sup>+</sup>-nuclei, MAP2<sup>+</sup>-neurons, and cleaved caspase 3<sup>+</sup>-apoptotic cells. Mean ± SEM, n = 3-6 wells per group from 1 differentiation. Statistical analysis: one-way ANOVA, \*p < 0.05.

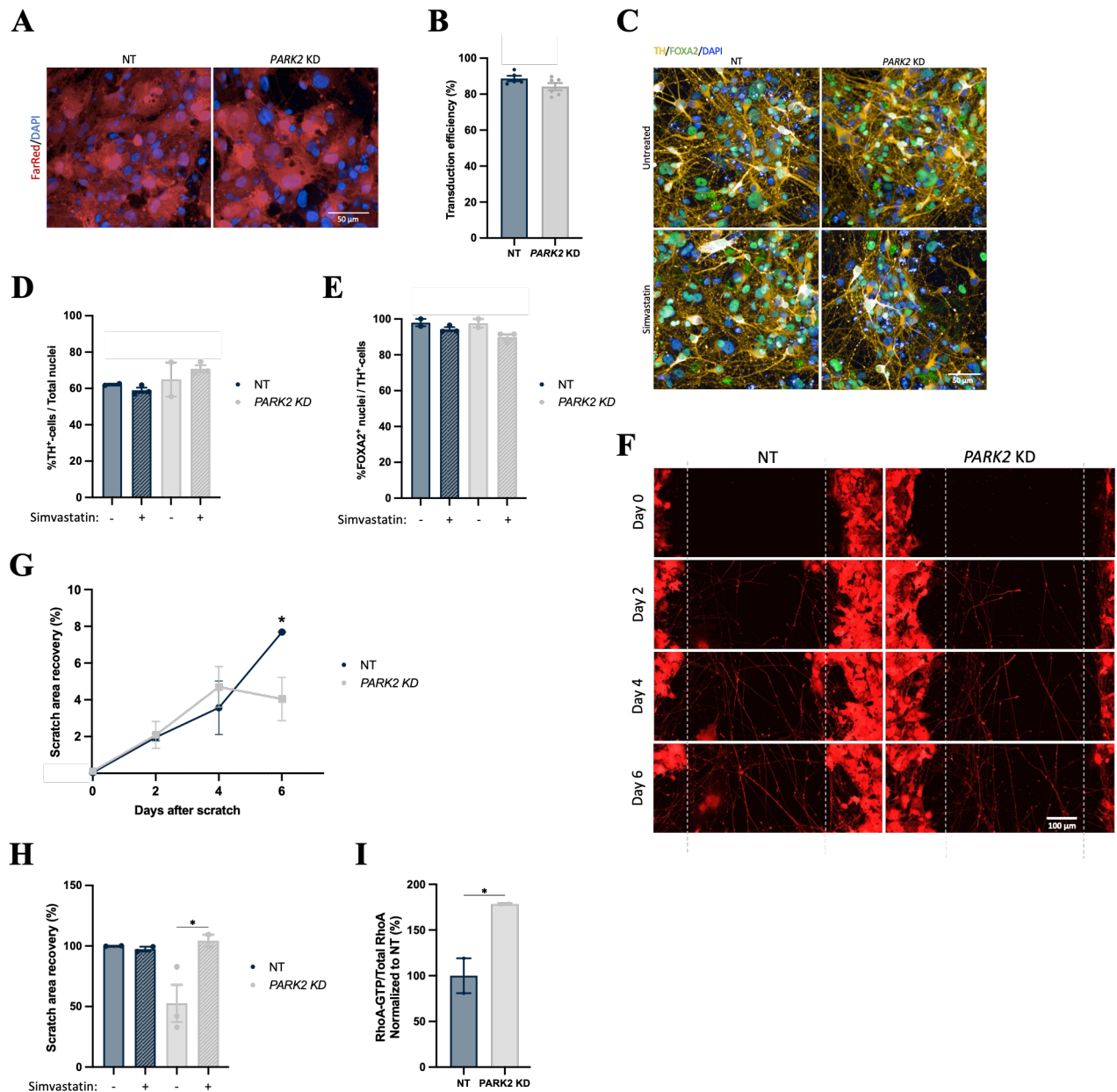

**Suppl. figure 9. CRISPRi knockdown (KD) of *PARK2*.** A-B) Transduction efficiency of CRISPRi-mediated KD of parkin in the differentiated healthy WTC11 iPSC line was quantified by the number of cells expressing the FarRed reporter gene. Mean ± SEM, n = 6 wells per group from 1 differentiation. Statistical analysis: Student's t-test. Scale bar: 50 μm. C-E) Immunofluorescence staining and quantification of tyrosine hydroxylase- (TH, yellow) and FOXA2- (green) expressing DA neurons, showing no effect of KD on the differentiation capacity of the cells. Simvastatin treatment was also found not to change the content of DA neurons in the cultures. Mean ± SEM, n = 2-3 wells per group from 1 differentiation. Statistical analysis: one-way ANOVA. Scale bar: 50 μm.

F-G) Scratch analysis of transduced cells was performed by imaging for the FarRed reporter gene at day 2, 4, and 6 post scratch. Decreased neurite outgrowth was seen upon *PARK2* KD 6 days post scratch compared to NT control cultures. Statistical analysis: Student's t-test comparing *PARK2* KD and NT for each timepoint, \*$p < 0.05$. H) Simvastatin treatment improved neurite outgrowth for *PARK2* KD DA cultures. Mean ± SEM, $n = 2-3$ wells per group from 1 differentiation. Statistical analysis: one-way ANOVA, \*$p < 0.05$. I) RhoA activity was upregulated upon *PARK2* KD. Mean ± SEM, $n = 2$ samples from 1 differentiation. Statistical analysis: Students t-test, \*$p < 0.05$.
